# Regulation of mesenchymal stem cell function by TGFβ-1 on mast cell extracellular vesicles – role of endosomal retention

**DOI:** 10.1101/172213

**Authors:** Yanan Yin, Ganesh Vilas Shelke, Su Chul Jang, Cecilia Lässer, Stefan Wennmalm, Hans Jürgen Hoffmann, Jonas Nilsson, Li Li, Yong Song Gho, Jan Lötvall

**Affiliations:** Krefting Research Centre, Institute of Medicine at the Sahlgrenska Academy, University of Gothenburg, Göteborg, Sweden.; Laboratory Medicine, Shanghai First Peoples Hospital, Shanghai, China.; Royal Institute of Technology-KTH, Department of Applied Physics, Experimental Biomolecular Physics Group, SciLife Laboratory, 171 65, Solna, Sweden.; Department of Respiratory Diseases and Allergy, Aarhus University Hospital.; Department of Clinical Medicine, Aarhus University.; Department of Surgery, Institute of Clinical Sciences, University of Gothenburg, Gothenburg, Sweden.; Department of Life Sciences, Pohang University of Science and Technology, Pohang, Gyeongbuk 790-784, Republic of Korea.; Codiak BioSciences Inc., 500 Technology Square, 9th floor, Cambridge, MA 02139, USA.

**Keywords:** Extracellular vesicles, exosomes, mesenchymal stem cells, tumor growth factor beta-1, cellular localization, lysosomal evasion

## Abstract

Extracellular vesicles (EVs) convey biological messages between cells, either by surface-to-surface interaction, or by shuttling of bioactive molecules to a recipient cell cytoplasm. Here we show that EVs released by human primary mast cells or transformed human mast cells (HMC1), carry TGFβ-1 on their surface. EV-associated TGFβ-1 enhance the migratory activity of human mesenchymal stem cells (MSCs) compared to free TGFβ-1, as both knockdown of TGFβ, or a TGFβ-antibody, attenuate the effect. The MSCs respond by increasing matrix metalloproteinase-2 and −9 (MMP) activity. Further, EVs given to MSCs are retained in the endosomal compartments at a time of biological function, prolonging EV-associated TGFβ-1 signaling vs free TGFβ-1. When exposed to EVs, MSCs home more toward allergen-exposed lung in a mouse allergen model, resulting in attenuated allergic inflammation. Our results show that mast cell-EVs are decorated with TGFb-1, are retained in endosomes, which influences both MSC phenotype and function.

## INTRODUCTION

Inflammatory responses involve intercellular communication through secreted soluble mediators such as chemokines and cytokines (*1-4*). These molecules induce different signaling pathways in cells, and modulate gene expression. Beyond soluble mediators, cells also release extracellular vesicles (EVs) that can influence recipient cell phenotype. EVs are nanometer-sized, lipid bilayer-enclosed membrane structures that carry an array of bioactive molecules, including proteins, nucleic acids, and lipids (*5-12*). EVs can rapidly be taken up by recipient cells, and the contents of the EVs can induce a vast array of phenotypic changes in a recipient cells, which has been implied in both health and disease (*13-16*).We and others have shown that EVs carry bioactive molecules on their surface, including c-KIT, Wnt and IL-1β (*6, 17*).

It has been shown that mesenchymal stem cells (MSCs) can regulate immune responses (*18-22*), which has made them attractive therapeutic candidates in several inflammatory diseases, including multiple sclerosis, rheumatoid diseases, tissue transplant rejection and inflammatory respiratory disorders (*23-25*). It is known that cytokines can alter the MSC phenotype in numerous ways (*26-29*), but very little is known whether or how EVs can influence MSC functions, and exactly by which mechanism this may be induced.

In theory, MSC biology *in vivo* may be regulated by cells in their vicinity, and thus the EVs released by other cells and taken up by the MSCs within the body have the capacity to regulate MSCs. Further, it could also be proposed that EVs may be used to manipulate MSCs *in vitro* to improve their therapeutic potential, putatively increasing their immunomodulatory effects as cell therapies in disease. We here hypothesize that mast cell-derived EVs influence the phenotype of human MSCs, and thereby their therapeutic potential in a model of respiratory inflammation. To verify this, we have utilized EVs from a both primary human mast cells as well as a human mast cell line, and tested the ability of EVs to influence the phenotype of primary bone marrow derived mesenchymal stem cells, including identifying intracellular signaling pathways that may be involved in an induced response. Further, we could identify a molecular component of EVs that can influence recipient cell biology. Lastly, the effects of EV-treated MSCs on a lung allergic inflammation model are tested.

## RESULTS

### Mast cell-derived EVs enhance the migration of MSCs in-vitro

EVs were isolated from the human mast cell line, HMC-1.2 by using differential ultracentrifugation. The mast cell-derived EVs had a size of 125.4 ± 3.9 nm according to nanoparticle tracking analysis (NTA, Supplementary Fig. 1a), and were positive for EV-enriched proteins such as CD63, CD81, and CD9 as detected by flow cytometry (Supplementary Fig. 1b). EVs from primary human derived mast cells, (cultured from a human peripheral blood stem cell population positive for CD133^+^) have similar size and structure on EM (40-120 nm) (Supplementary Fig. 1c), and express CD63 as detected by flow cytometry (Supplementary Fig. 1d). HMC-1 EVs were efficiently taken up by primary human MSCs in a time and temperature-dependent manner, which suggests that the EV uptake is a biologically active process (Supplementary Fig. 2a-c). Adding EVs to the cell cultures induced a n elongated morphology of the MSCs (Fig. 1a), but did not alter the ability of the MSCs to differentiate into adipocytes or osteocytes (Supplementary Fig. 4). *In vitro* analysis of an MSC scratch assay, suggest that the EVs increase wound healing activity (Fig. 1b). Enhanced migratory activity of MSCs upon EV stimulation was also associated with increase in transcripts of matrix metalloproteinases (MMP-2 and MMP-9) mRNA in cells, and with increased secretion of MMP-2 and MMP-9 proteins into the cell culture supernatant, associated with increased gelatinolytic activity (Fig. 1c-d). It has previously been shown that multiple growth factors can influence MSC migration behavior by regulating signaling cascades(*30*). We therefore exploited fetal bovine serum (FBS) as a pan-chemoattractant in the next experiment, and could show that EV-treated MSCs have a higher migratory tendency towards the FBS compared to non-treated MSCs, in a dose-depended manner (Fig. 1e). Collectively, these results show unequivocally that mast cell-derived EVs enhance the migration of MSCs *in vitro*.

**Figure 1: Mast cell-derived EVs enhance the migration and invasion of primary human mesenchymal stem cells.**
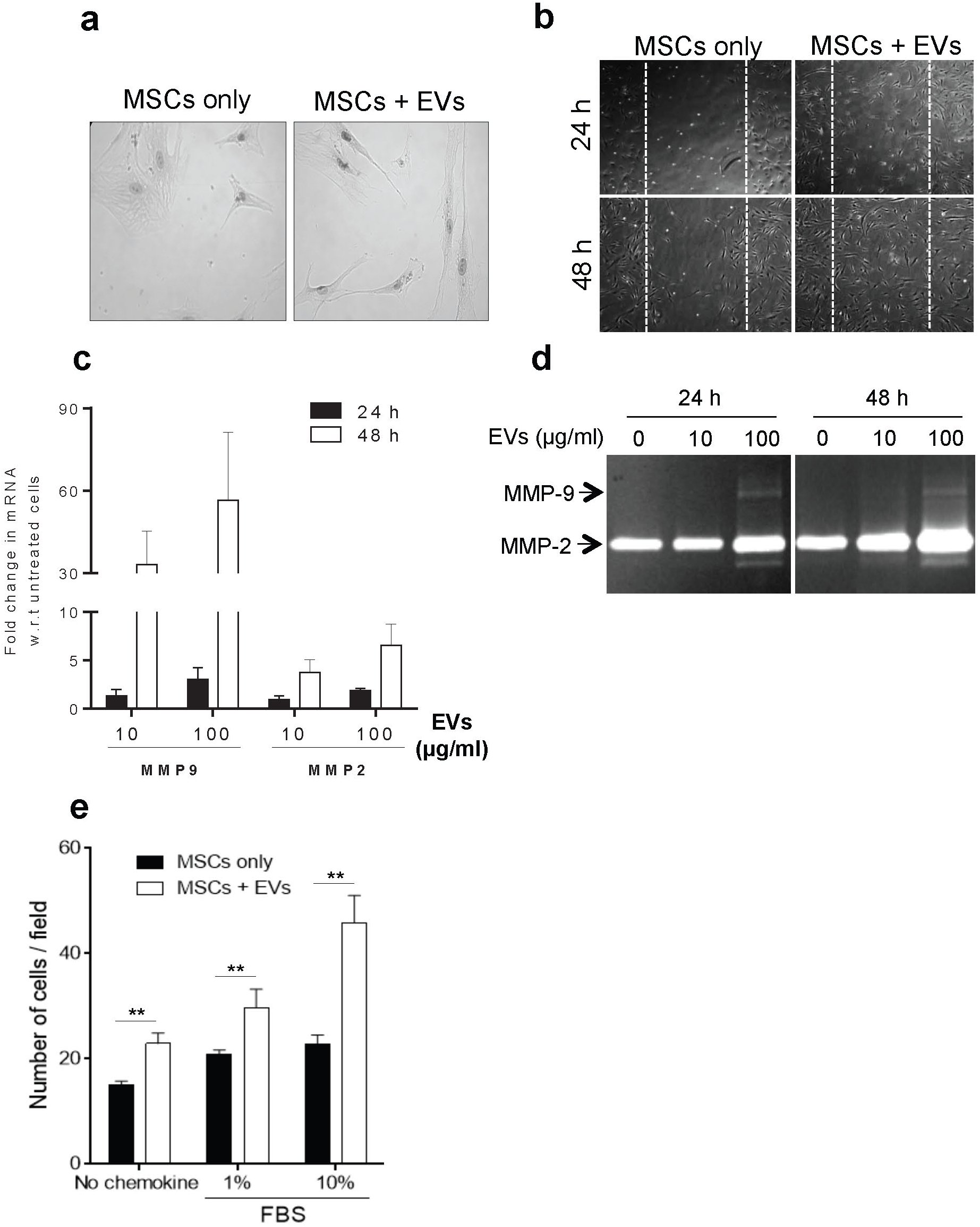
**(a)** EVs induce morphological changes in MSCs as determined by Giemsa staining. (**b**) An *in vitro* wound healing scratch assay was performed on monolayers of untreated or EV-treated MSCs (n = 2). Plates were imaged at 24 h and 48 h after injury. **(c)** MSC mRNA transcripts of MMP-9 and MMP-2 after EV treatment as determined by PCR. W.r.t; with respect to. **(d)** Gelatinolytic activity in secreted supernatant from MSCs estimated by zymography to detect MMP-2 and MMP-9 activity. **(e)** Invasiveness of EV-treated MSCs towards different concentrations of FBS evaluated using a Boyden chamber Matrigel assay. Data are presented as means ± SEM; n = 4; ** p ≤ 0.01.

### TGFβ-1 is present on the surface of mast cell-derived EVs

To identify surface components of mast cell-derived EVs that could potentially regulate the observed MSC migration, we first determined the membrane proteome of the EVs. This proteomics approach reduces extra and intra-vesicular proteins, but enriches membrane proteins and therefore identifies various low-abundance bioactive proteins that are not detected in standard EV proteomics assays (*31*). In total we identified in total 1743 proteins (Supplementary Table 1), of which 504 were membrane-associated proteins, including several receptors such as TGFβ and insulin receptors. Interestingly, we also detected cytokines such as TGFβ-1 that are rarely identified with mass spectrometry due to their low abundance. Multiple approaches were used to validate the presence of TGFβ-1 on the EVs. Firstly, we developed a classical sandwich ELISA to detected the relative abundance of TGFβ-1 in TGFβ-1+ as well as CD63+ EVs in various fractions of HMC-1 EVs after iodixanol density gradient separation. Confirming our proteomics finding we determined that the majority of TGFβ-1floated in fractions no. 1 and 2, with density in the range of 1-1.08 g/ml (Supplementary Fig 3 and Fig. 2a. This was coincident with the EV markers TSG101 and CD81. In addition, the association of TGFβ-1 with EVs was observed by fluorescence correlation spectroscopy (FCS). Specifically, we found that the signals for TGFβ-1 and the EV marker CD63 were co-localized in the same time lapse, and had the same diffusion time in the FCS analysis (Fig. 2b), which strongly suggests that they are co-localized on the same individual EV. With this method, we determined that approximately 17 % of CD63 positive EVs were also positive for surface expression of TGFβ-1, but some EVs carried either marker alone. Finally, we performed a sandwich ELISA to quantify the amount of TGFβ-1 associated with vesicles bound to anti-CD63-coated beads. We could detect approximately 12 pg of total TGFβ-1 on 30 µg CD63-positive EVs from HMC-1 cells, out of which 7.1pg was of the active form of TGFβ-1 (Fig. 2c). Taken together these experiments conclusively show that TGFβ-1 is associated with EVs on their surface.

**Figure 2: Bioactive TGFβ-1 co-localizes with EVs.**
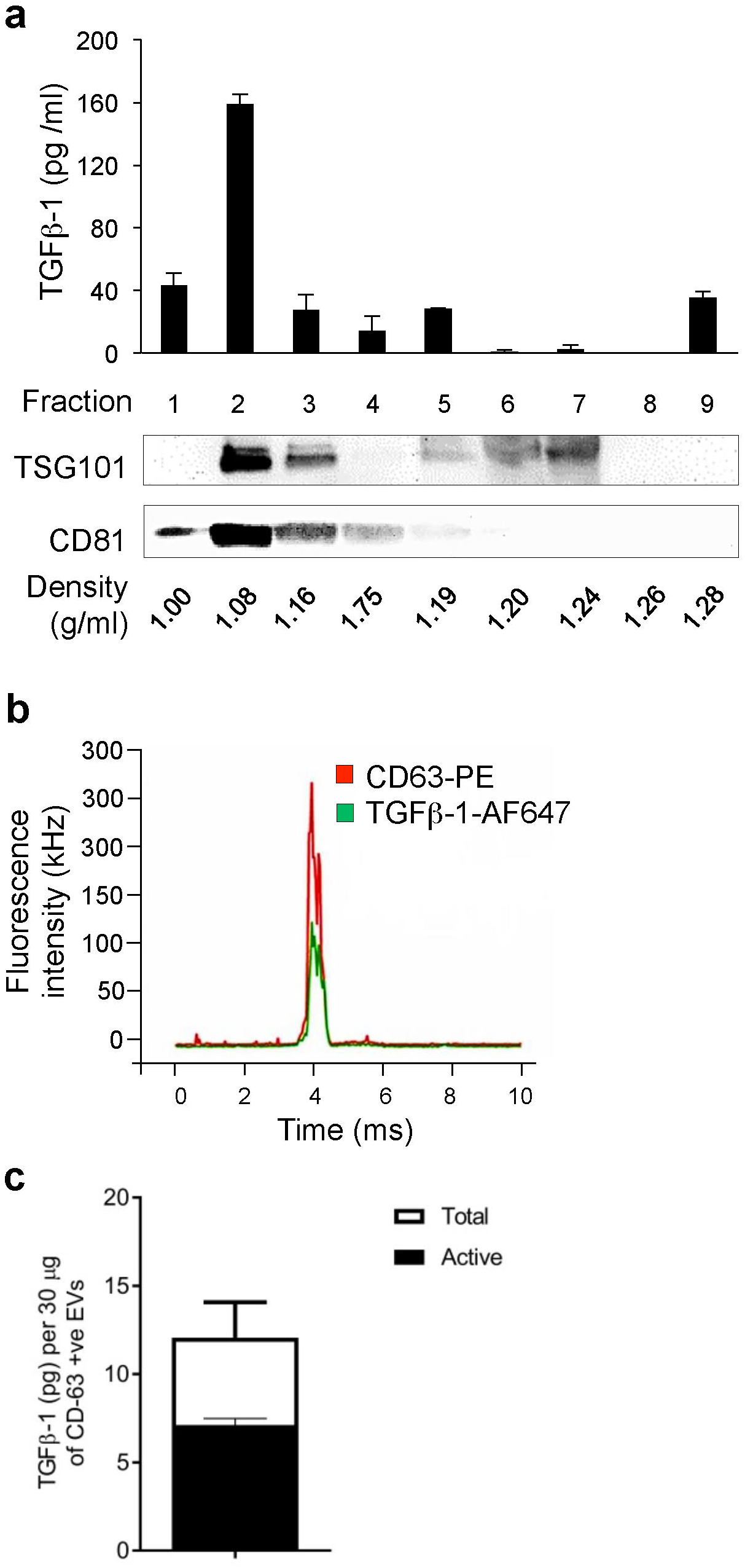
(**a**) EVs isolated from mast cells were floated on OptiPrep gradients, and the expression of TGFβ-1 (ELISA) and the EV markers TSG101 and CD81 (immunoblotting) were measured. **(b)** Quantitative measurements of the distribution of TGFβ-1 and CD63 on single EVs (DiO labeled) using fluorescence correlation spectroscopy (FCS). Excitation was at both 488 nm and 594 nm. The width of the vesicle, more than 500 ms, corresponds to a vesicle diameter of several micrometers. **(c)** The inactive and active forms of TGFβ-1 on the EVs were measured per unit of CD63-positive EVs using ELISA. Data are presented as means ± SEM (n = 3).

### TGFβ-1 on the surface of EVs induces enhanced migration of MSCs

As it has been shown before that free, soluble TGFβ-1 can stimulate MSC migration (*27*) and since we in the proteomics observed TGFβ-1is present in EV membrane isolates, we set out to determine whether this molecule is responsible for the observed EV-mediated enhanced migration of the cells. Importantly, SMADs are a family of proteins that are involved in signal transduction and transcriptional regulation (*32*), and SMAD2 is activated in response to TGFβ signaling (Fig. 3a) and could be involved in MSC migration (*33*). Indeed, we observed that the ratio between nuclear and cytoplasmic SMAD2 in MSCs increases following EV exposure, indicating that EVs induce nuclear translocation of this transcription factor (Fig. 3c). Consistent with activation of SMAD2, we also observed an increase in *Tgfb1* mRNA transcript following stimulation with EVs, in a time- and dose-dependent manner. This suggests that EVs enhance *Tgfb1* expression in MSC, most likely via SMAD2 signaling (Fig. 3d). This is consistent a previous study documenting that soluble TGFβ-1 regulates its own expression in an autocrine manner (*34*).

**Figure 3:**
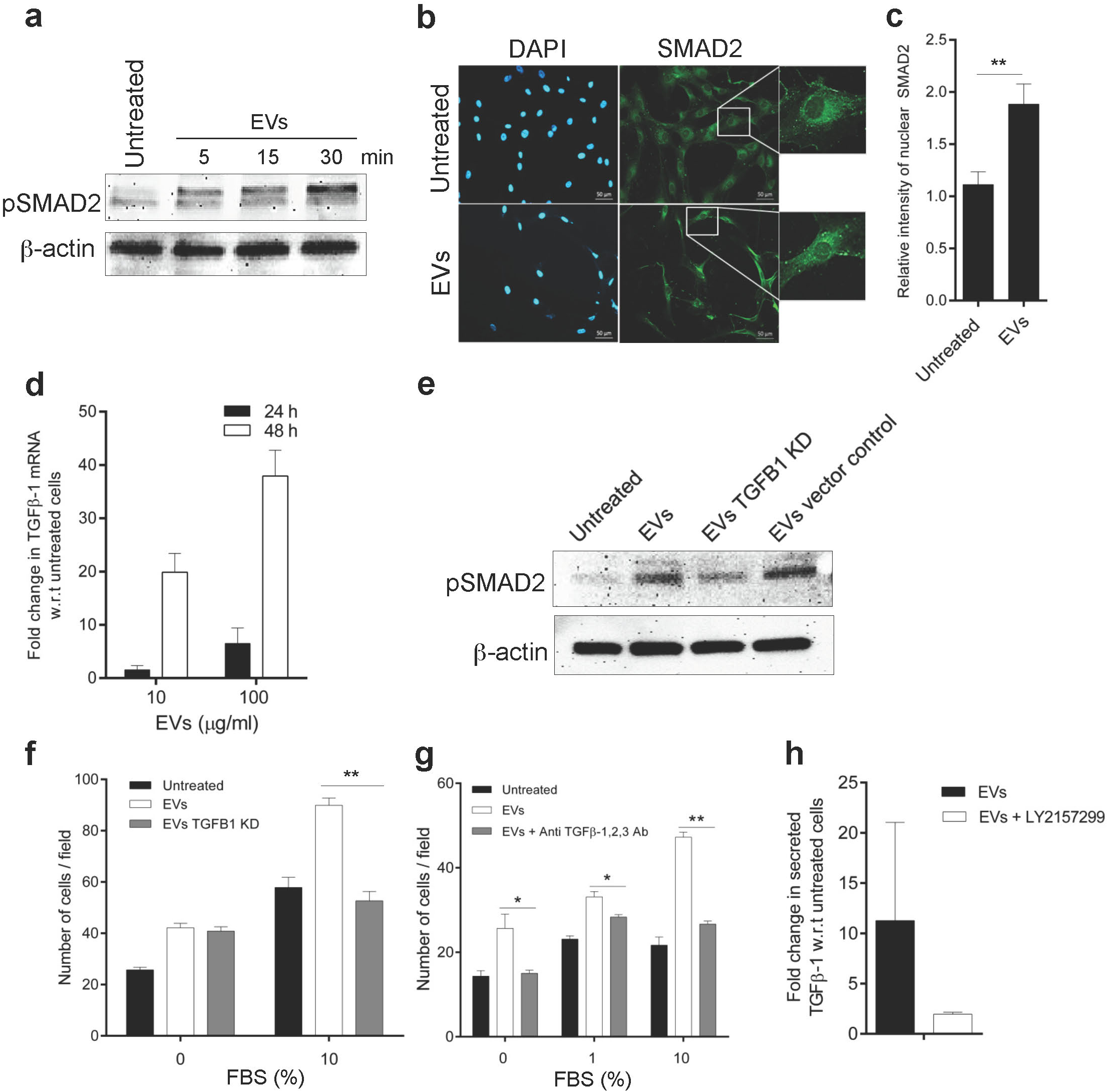
EVs activate SMAD signaling in human mesenchymal stem cells. **(a)** EVs (100 µg/ml) activate the TGFβ-1 signaling axis by phosphorylation of SMAD as identified by immunoblotting at different time points. **(b, c)** Activation can be visualized by immuno-fluorescent imaging as the translocation of total-SMAD in the nucleus (**b**) and can be quantified by the relative intensity measurement (c) in MSCs 30 min after EV exposure. **(d)** Activation of transcripts downstream of SMAD, such as TGFB1, was evaluated in MSCs at 24 h and 48 h after EV exposure. **(e, f)** EVs from TGFβ-1 knockdown HMC-1.2 cells were used to induce phosphorylation of SMAD in MSCs as measured using immunoblotting **(e)** and as measured by their capacity to induce migration towards FBS **(f)**. **(g)** Invasiveness of MSCs towards FBS after EV treatment was also evaluated in the presence of a blocking antibody against TGF-β-1, 2, 3 using a Boyden chamber invasion assay. **(h)** mRNA levels of TGFβ-1 in MSCs were measured in the presence of the TGFR-I receptor blocking agent LY2157299 during exposure to EVs. Data are presented as means ± SEM; n = 3; * p ≤ 0.05, ** p ≤ 0.01.

To further determine whether the EV-associated TGFβ-1 is responsible for the observed enhanced migration and TGFβ-1signaling in MSCs we first isolated EVs from TGFB1 knock-down (*35*) HMC-1 cells, using a doxycycline-inducible CRISPR/Cas-9 system (Supplementary Fig. 6a-b). Indeed, EVs derived from TGFβ-1 KD mast cells resulted in reduced phosphorylation of SMAD2 in recipient MSCs compared to controls (Fig. 3e), as well as reduced migration towards the pan-chemoattractant FBS (Fig. 3f). To further confirm this, we pre-incubated EVs with a TGFβ-neutralizing antibody that blocks the TGFβ-1 interaction with its receptor. Our results show that TGFβ-neutralized EVs evoke less phosphorylation of SMAD2 (Supplementary Fig.6) and significant reduction in the migration of the MSCs towards the pan-chemoattractant FBS (Fig. 3g). In addition, we found a drastic drop in TGFβ-1 secretion into the growth media by the MSCs when the TGFβ-1 receptor was blocked by a specific small molecule inhibitor (LY215799) (Fig. 3h). Collectively, these results suggest that TGFβ-1 on the EV surface induces an enhanced migration phenotype and TGFβ-1 response in MSCs.

### Sustained TGFβ-1 signaling by endosomal retention of EVs and lysosome evasion

It is well known that TGFβ-1 rapidly activates signaling by phosphorylation of SMADs, which then quickly lessens (*36, 37*). Having observed similar signaling activation and functional outcome for EV-bound TGFβ-1, we asked whether free TGFβ-1 induces similar temporal signaling activation in MSCs in our system. To test this, we used dose-matched concentrations of TGFβ-1 in its free form and in the EV-bound form, and performed Western blotting analysis for phosphorylated SMAD2 (pSMAD2) in the recipient MSCs. First, we determined the amount of TGFβ-1 associated with our total EV-isolates after floating them on a density gradient. We found approximately 18.5 pg of total TGFβ-1 in 30 µg total EVs, out of which approximately 40% was in the active form of TGFβ-1 (Supplementary Fig.7). Hence, 0.3 pg of active TGFβ-1 was considered to be present per microgram of total EVs, which was utilized for further experiments (Supplementary Fig. 7). Interestingly, TGFβ-1 displayed altered activation kinetics when bound to EVs, producing a lower amplitude (Supplementary Fig. 8 (30 mins)), but more sustained (Fig. 4a (60 mins)) activation of pSMAD2. We further found that EVs stimulate MSC migration more efficiently than free TGFβ-1 at dose-matched concentrations (≥30 pg/ml) (Fig. 4b). These data suggest that the sustained, low amplitude signaling elicited by EV-bound TFGβ-1 has more functional effect than the transient, high amplitude signal evoked by free TFGβ-1.

**Figure 4:**
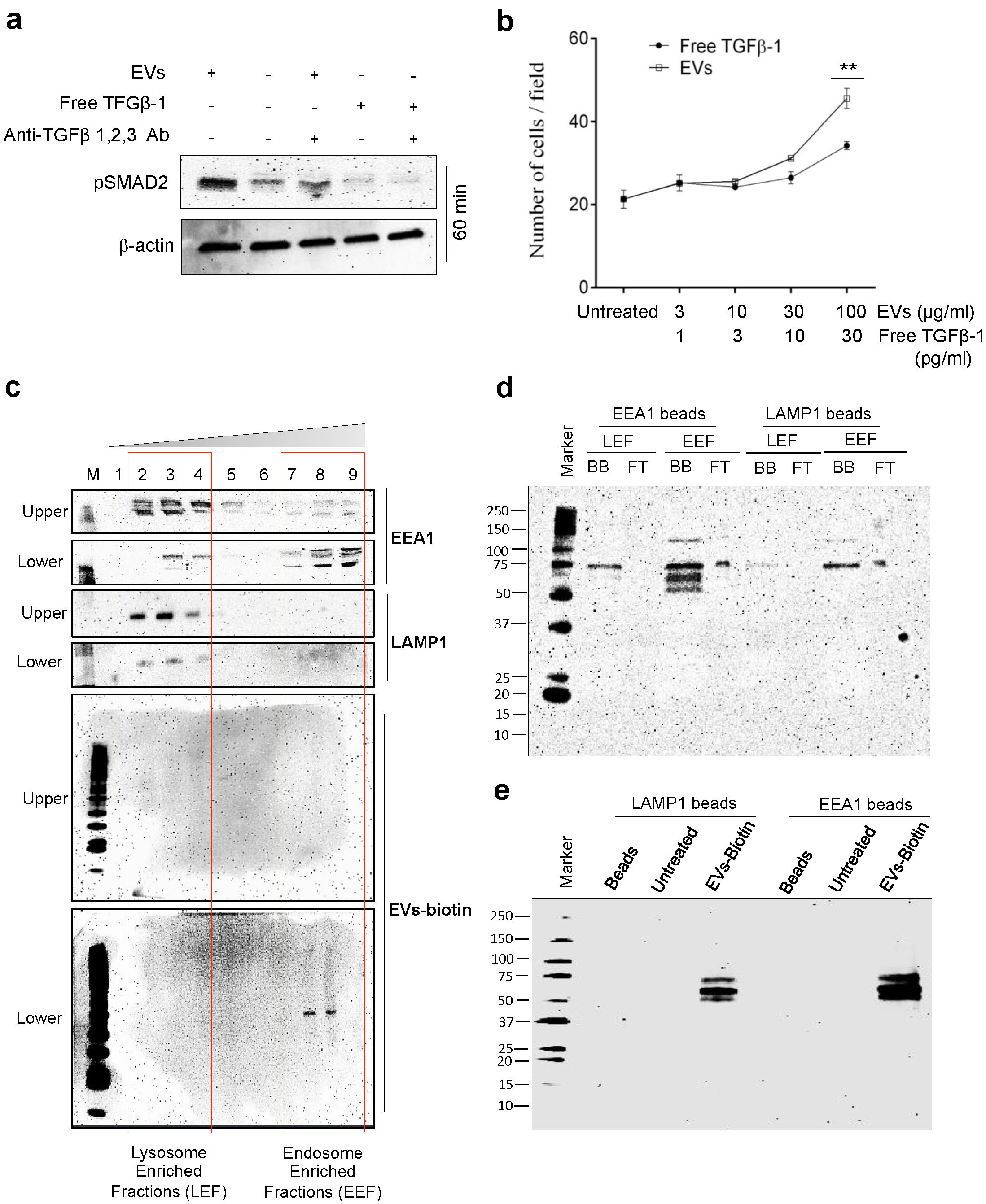
TGFβ-1 associated with EVs has higher signaling stability compared with free TGFβ-1. (**a**) The signaling efficiency in MSCs, as determined by immunoblotting of pSMAD, in response to dose-matched concentrations (10 pg/ml) of free TGFβ-1 or EV-bound TGFβ-1 estimated at 60 min. **(b)** The migration of MSCs towards free TGFβ-1 or EV-bound TGFβ-1 as a chemoattractant was performed by reverse migration assay. Data are presented as means ± SEM; n = 3; * p ≤ 0.05. **(c)** Biotinylated EVs were incubated with HEK293-T cells for 60 min, and organelles were isolated with optiprep gradient as described in Supplementary Fig. 9 and probed to detect LAMP1, EEA1 and biotinylated EVs distribution. **(d)** Organelles isolated with optiprep gradient from HEK293-T treated with biotinylated EVs followed by LAMP1 or EEA1 antibody based capture and were probed for biotinylated EV proteins. **(e)** Presence of biotinylated EVs at 60 mins were detected in crude cytoplasmic preparation of MSCs cells that were bounded to LAMP1 or EEA1 antibody beads coated.

To elucidate the difference in the signaling efficiency of free versus EV-bound TGFβ-1, we assessed the fate of EVs in intracellular compartments, primarily comparing lysosomes and early endosomes. It has previously been reported that free TGFβ-1 is rapidly processed via lysosomes (*38*), and we hypothesized that EV-bound TFGβ-1 taken up by recipient cells evades or is delayed in trafficking to lysosomes, producing a prolonged TGFβ intracellular signal. As it is not feasible to perform these experiments in large scale using MSCs, we performed these specific organelle localization experiments on HEK-293T cells. Briefly, we performed high-resolution fractionation of cell organelles, to physically separate lysosomes and early endosomes from HEK-293T cells that were exposed to biotinylated-EVs for 60 min (Supplementary Fig. 9a and Fig. 4c). We found that the endosome-enriched fraction (EEF) was highly enriched in biotinylated proteins as compared with the lysosome-enriched fraction (LEF), suggesting that the EV components taken up by the cells preferentially are retained in endosomes, where they may evade lysosomal degradation at the time of activating the cell (Fig. 4c). The observed prolonged EV-TGFβ-1 mediated activation of cells may therefore at least partly be explained by EV lysosomal evasion.

To confirm the endosomal localization of the EV components, two different approaches were taken. First, EEF and LEF samples were incubated with organelle-specific antibodies to pull down the endosomal and lysosomal fractions (Fig. 4d), again showing enrichment of biotinylated proteins in the EEA1 specific pull-down (as a marker of endosomes) compared to the LAMP1-specific pull-down (as a marker of lysosomes) (Fig. 4f). To confirm this finding in human MSCs we traced the biotinylated EVs by performing a similar organelle specific pull-down of EEA1 and LAMP1 from crude cytoplasmic preparation of MSCs. As for the HEK-293T cells, also MSCs showed a higher biotinalylation signal in the endosomal compartments compared to the lysosomal compartments. Secondly, we looked for endosomal traces of streptavidin captured EV-biotin to validate the finding. Cells were incubated with biotinylated EVs and then we used streptavidin-coated beads to pull down the cellular compartments containing biotinylated proteins where the EV-associated biotinylated proteins are present, as described in Supplementary Fig. 9b (Left panels). As expected, the total amounts of proteins binding to the beads were significantly higher in the EEF compared to the LEF sample (Supplementary Fig. 9b (i)). Furthermore, a higher percentage of the proteins were bound to the streptavidin-coated beads in the EEF samples, while a higher percent of the proteins were unbound to the beads in the LEF sample, indicating that more biotinylated proteins are present in the endosome enriched fraction. To further confirm that the bead-bound material (biotinylated proteins) is originating from the organelles of interest, we identified the early endosomal protein, EEA1 (here used as a marker for the early endosome) and LAMP1 (here used as a marker for the lysosome) (Supplementary Fig. 9b (ii)) in the bead-bound samples. In summary, the results of these collective experiments strongly argue that the EVs remain longer in the early endosomes, suggesting early avoidance of traffic of the EVs to the lysosomes, thus allowing for prolonged TGFβ-1 signaling in the recipient cell.

### Functional enhancement of MSCs upon EV treatment in protecting allergic inflammation

To detect changes in the *in vivo* function of MSCs induced by EVs, we used a mouse model of ovalbumin (OVA)-induced lung inflammation treated with MSCs. The MSCs were engineered to express luciferase and green fluorescent protein (Luc-GFP-MSCs). Luc-GFP-MSCs were cultured with or without the introduction of HMC-1.2-derived EVs prior to being injected intravenously into the tail vein of OVA-sensitized and allergen challenged mice (Supplementary Fig. 10). The distribution of the MSC-derived luciferase activity in the mice was examined at four different time points (10, 30, 60 and 120 mins). The injected MSCs primarily localized to the lung, but administration of EV-treated Luc-GFP-MSCs resulted in relatively higher luciferase signal in lungs compared to the untreated Luc-GFP-MSCs, especially at the early time points (Fig. 5a). Moreover, we detected increased number of EV-treated Luc-GFP-MSCs in the lung parenchyma using fluorescence microscopy, compared to non-treated Luc-GFP-MSC, as seen by the detected GFP signal at 72 h post injection (Fig. 5b). We could also observe that EV-treated MSCs significantly reduced eosinophilia in the bronchiolar alveolar lavage fluid (BALF) in OVA-exposed mice, as compared to mice given the same number of untreated MSCs (Fig. 5c). These results support the hypothesis that mast cell-derived EVs enhance the migratory activity of MSCs to the inflamed lung, which results in an immune-regulatory function resulting in reduced allergic inflammation *in vivo*.

**Figure 5:**
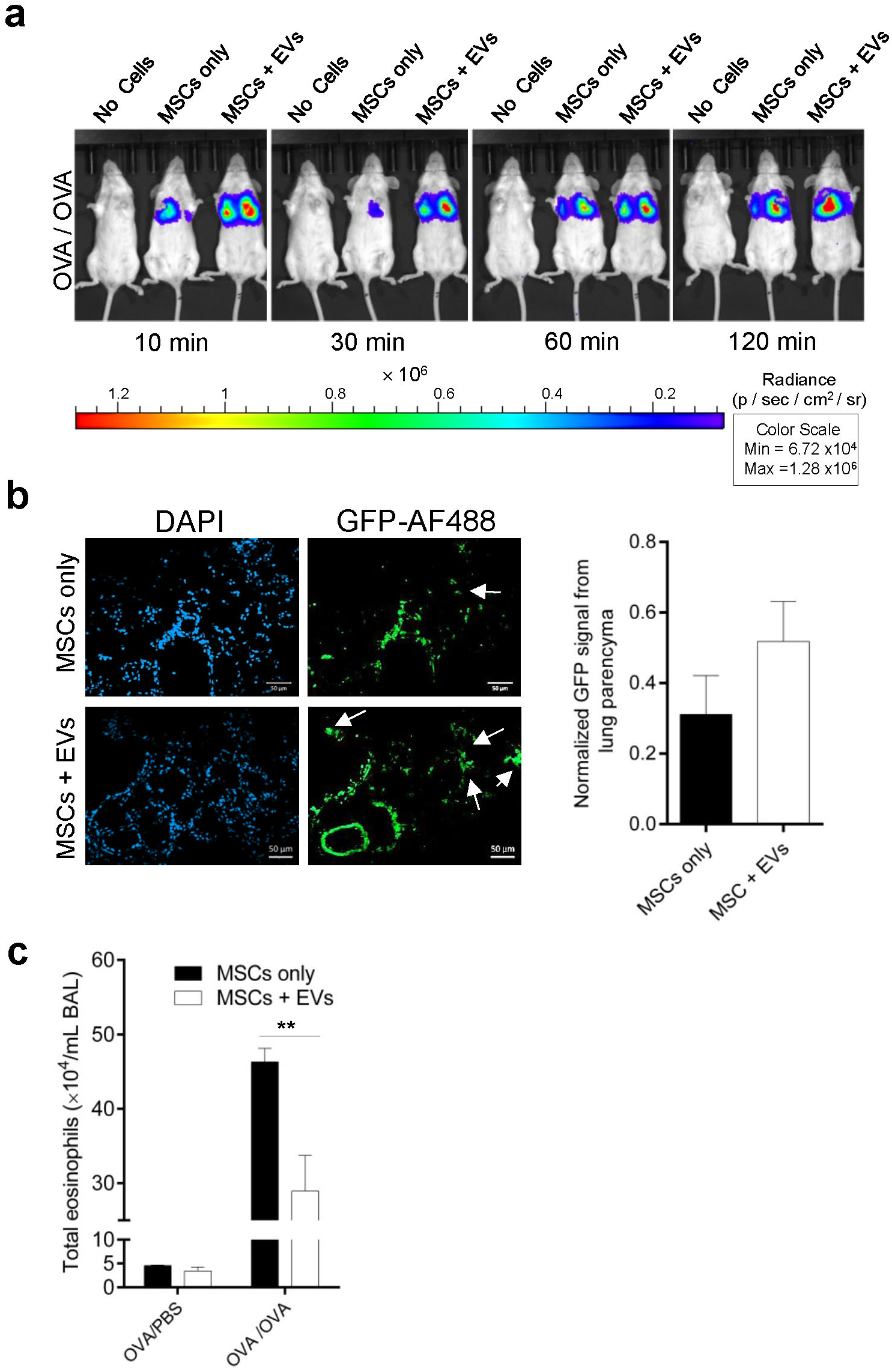
Mast cell-derived EVs enhance anti-inflammatory properties of primary human mesenchymal stem cells in allergic inflammation. **(a)** MSCs constitutively expressing luciferase were incubated with EVs *in vitro* for 48 hours prior to the MSCs being injected into the tail vein of OVA-sensitized and challenged mice (*in vivo* model of inflammation of the lung). The bioluminescence (photons·s^−1^ ·cm^−2^) measurement was performed in the OVA/OVA mice at 10 min, 30 min, 60 min, and 120 min after receiving the EV-treated or untreated Luc-GFP-MSCs. **(b)** Luc-GFP-MSCs in lungs tissue were stained with an AF488 labeled antibody against GFP and DAPI (Bar = 50 μm) and signal intensity was measured. **(c)** Eosinophils were counted in bronchiolar alveolar lavage fluid (BALF) to measure the degree of inflammation in the lung (n = 3-5). Data are presented as means ± SEM; ** p ≤ 0.01.

## DISCUSSION

Human MSCs have the capacity to regulate multiple types of inflammation, and are being tested clinically in multiple inflammatory diseases. We here show that mast cell-derived EVs in a dose-dependent manner induce an enhanced migratory MSC phenotype, and this effect is associated to SMAD2 signaling in the cell. This response is induced by TGFβ-1 present on the surface of the mast cell EVs, as shown by both TGFβ-1 antibody blocking experiments, as well as fluorescent correlation spectroscopy with single-EV resolution. The biological effect of the EV-associated TGFβ-1 is more potent and extended vs. that of free TGFβ-1. We further observed that EVs traffic primarily the endosomal compartment of the recipient cells, and less so to the lysosomal compartment, at a time point when SMAD2 phosphorylation was significant. Lastly, the MSCs treated with mast cell EVs had a greater tendency to home to allergen-exposed mouse lung than untreated MSCs, and this increased homing was associated with reduced eosinophilic inflammation in the lung.

In response to injury or inflammation, MSCs have been reported to provide repair and immune-suppressive functions, which putatively could be further enhanced by any local inflammation (*39*). We here discovered that mast cell derived EVs change the phenotype of MSCs to a more migratory phenotype, which could alter their functionality at a site of inflammation. The increased migration of the MSCs was associated with increased expression of both matrix metalloproteinase 2 and 6, which also suggests that the cells could become not only migratory, but also more invasive. This was indeed confirmed by a Matrigel invasion assay, in which the MSCs were dose-dependently more migratory. We have previously suggested that bioactive molecules on EV surfaces may influence recipient cells, specifically the stem cell growth factor receptor c-kit, which may enhance recipient lung cancer cell growth (*6*). In the current study we utilized a stringent EV membrane proteomics approach to identify the low-abundance TGFβ-1 and its receptor (Supplementary Table 1). Validation of the membrane proteomics data, utilizing single-vesicle resolution, confirmed the presence of TGFβ-1 on the EV surfaces (Fig. 2b), which may suggest that this highly bioactive cytokine could mediate immediate biological responses when interacting with a cell, via TGFβ-1 receptor activation, partly explaining the observed change in MSC phenotype. Previous studies have shown that free signaling factors like PDGF, TGF-β, and FGF can induce growth and differentiation of MSCs and change the fate of MSC function, but our current study suggests that the cytokines actually are more potent if associated to the EV membrane. It is possible that this enhanced potency depends on the three dimensional association of the TGF-β-1, facilitating its interaction with its receptor on the recipient cell. Importantly, blocking EV-associated TGFβ-1 with an antibody was sufficient to significantly reduce TGFβ signaling, further supporting the importance of this bioactive molecule for the observed biological effects in MSCs.

A detailed evaluation of the EV-associated TGF-β-1 suggested that approximately 40% is in active vs 60% inactive forms of TGFβ-1 (Supplementary Fig. 7). This distribution of TGFβ-1 was different from that seen in a previous study by Webber et al (2010) where they suggested that only ~2% of the TGFβ-1 from mesothelioma or prostate cancer cell line-derived EVs was in active form (*40*). This further emphasizes that mast cell-derived EVs may have a much higher proportion of active TGFβ-1, than EVs derived from other cells, which potentially could favor mast cell EVs to influence MSC function, putatively to enhance the therapeutic effects of these cells.

Importantly, it has been shown that free TGFβ-1 and EVs from tumor cells (prostate and breast cancer cells) can induce differentiation of MSCs into osteogenic and myogenic lineages respectively (*41, 42*). The effects of free TGFβ-1 in driving differentiation into cellular lineages has been argued to take place in a context-dependent manner, and that other niche-specific may be important to direct the MSC fate (*43*). This feature of commitment to differentiation is not desirable in the clinical setting if MSCs are being administered to patients, as they could give rise to unwanted mesenchymal or otherwise harmful phenotypes. Importantly, using linage specific medium for adipocytes and osteocytes, we could demonstrate that MSCs treated with TGF-β-1 positive EVs retain multipotency, despite the observed phenotypic changes induced by the TGF-β-1-EVs. It is important to note that TGFβ-1present on the EVs was not able to induce lineage commitment in the MSCs, despite the previously reported effects of free TGF-β-1, which may suggest that that other cargo components in the EVs such as for example IL-9, insulin and TNF receptor (all which were identified in our membrane proteomics), could have a counteracting role in attenuating an undesirable differentiation of MSCs (*40*).

In all biological systems free biomolecules such as cytokines interacts with surface receptor on cells, resulting in downstream cellular signaling (*44, 45*). It may be counterintuitive to observe that the low dosages (picograms) of TGFβ-1 on the surface of EVs could contribute so significantly to downstream signaling, for example phosphorylation of SMAD2 (Fig. 3e). It is possible that exosomes present molecules to recipient cells in a very unique context, which could explain the increased potency of EV-associated TGFβ-1 compared to free molecules. For example, it has been shown that EVs enhance the oligomerization of prions proteins (non-toxic PrP^c^ to toxic PrP^sc^) by generating high local concentrations of template and PrP^c^ proteins and bringing them to close proximity (*44*). Thus, emerging evidence suggests that much lower concentrations of bioactive molecules can induce a stronger response when associated to the EVs.

When comparing the effects of EV-associated TGFβ-1 with free TGFβ-1, we could see both enhanced and prolonged effects of the EV-associated cytokine, both in relation to migratory function and pSMAD signaling in MSCs at a 60 minutes time point. It has previously been shown TGFβ-1 alone induces signaling by the activation of the TFGβ-1 receptor, and this complex is rapidly internalized and is transferred to lysosomes for degradation (*46, 47*). The fate of EVs in recipient cells, which is known to involve endosomal translocation (*48*) (*49*), could at least partially explain the longer signaling half-life of EV-bound TGFβ-1 over free TGFβ-1 observed in the current experiments. To investigate this, we isolated early endosomal and lysosomal compartments, and observed that EVs were preferentially located within the early endosomal compartment (EEA1-enriched), with a small amount of EV proteins in the lysosomal compartment (Fig. 4d-e)(*50*). Therefore, the presence of TGFβ-1 on the surface of EVs seems to allow its prolonged signaling, probably by avoiding degradation in the lysosomes. This could indeed be a novel strategy for long-term signaling by low abundance bioactive molecules, and putatively expression of other bioactive molecules on the surface of EVs could be important pathway by which EVs change recipient cell phenotype.

The increased migratory phenotype of mast cell-derived EV-treated MSCs *in vitro*, and that EVs enhance MSC homing towards inflammatory cues in mouse lungs *in vivo* that leads to a reduction in eosinophil recruitment, a signature of reduced lung inflammation. EV-treated MSCs may thus be useful for treating lung disorders in which inflammation plays a significant role, such as asthma (*39*), idiopathic pulmonary fibrosis (*51*), and acute lung injury (*52*). Previous approaches to treating different diseases with MSCs include pre-treatments of MSCs with cytokines such as TNF-α, IFN-γ, and IL-1β, and these have resulted in some success. However, these approaches have suffered from problems like differentiation, loss of immune-suppressive function, and limited target usage in certain treatments (*53*). Unlike other methods, MSCs pre-treated with EVs maintain their multi-potency (Supplementary Fig. 4) and immuno-suppressive effects, which makes them interesting therapeutic candidates in diverse diseases. We here observed an enhanced protective effect of MSCs pre-treated with EVs compared to untreated MSCs in an OVA-induced allergen inflammatory model (*54*). However, it is not yet clear if the enhancement of MSC function that we observed is because of early retention of MSCs in lungs, enhanced immunosuppression or both. Nevertheless, the *in vivo* imaging and MSC-derived GFP signal in the lung parenchyma suggests that both mechanisms operate simultaneously in reducing inflammation in the OVA model

Collectively, our findings suggest that EVs from mast cells significantly can change the phenotype of MSCs. In this study we have documented the presence of TGFβ-1 on the surface of a subset of mast cell-derived EVs, and this cytokine will regulates MSC. Also, the EVs remain in the endosomal compartment, and may in that way extend the biological function of the surface TGFβ-1. The present work also suggests that designing EV surface or cargo might be a useful novel strategy for manipulating MSCs, putatively to make them more potent therapeutic tools.

## METHODS

### Cell culture

Human bone marrow-derived mesenchymal stem cells (MSCs) were obtained at passage 1 from the MSC distribution at the Institute of Regenerative Medicine at Scott and White, USA. The MSCs were cultured in Minimum Essential Medium α-GlutaMAX™ Supplement (Life Technologies, Thermo Fisher Scientific, Waltham, MA, USA) supplemented with 15% fetal bovine serum (FBS; Sigma Aldrich, St. Louis, MO, USA). The culture media was changed to EV-depleted FBS-containing medium 24 hours prior to experiments. The MSCs were used within 3-4 passages in all experiments, with a seeding density of 3000 cells/cm^2^. The human mast cells, HMC-1.2 (J, Butterfield, Mayo Clinic, Rochester, MN, USA), were grown in Iscove’s modified Dulbecco’s medium (IMDM, HyClone Laboratories, Logan, UT, USA) supplemented with 10% EV-depleted FBS, 2 mM L-glutamine (HyClone Laboratories) and 1.2 mM α-thioglycerol (Sigma Aldrich). HEK-293T cells (ATCC, Manassas, VA, USA) were grown in RPMI-1640 medium (HyClone Laboratories) supplemented with 10% FBS. All cultures were supplemented with 100 units/ml penicillin and 100 µg/ml streptomycin (HyClone Laboratories). Here, the supplement FBS was depleted of EVs by ultracentrifugation for 18 hours at 120,000 × g (Type 45 Ti rotor, Beckman Coulter). All cells were cultured at 37°C in a 5% CO_2_ humidified atmosphere.

For the purification and culture of progenitor mast cells we used PBMC from health human donor. Briefly, mononuclear cells were purified from PBMC, and CD133^+^ cells population using MACS (Miltenyi Biotech, Germany). CD133^+^ cells were cultured in a serum free medium (StemSpan, StemCell technology, Vancouver Canada) supplemented with SCF and IL-6. IL-3 was added for first two weeks, and later it was removed. Cells then were maintained for 6-7 weeks before the conditioned medium was harvested for EVs isolation.

### Mice

B6-albino mice were purchased from Jackson Laboratories. Age matched females mice were used in the experiments at 6 to 10 weeks old. Pathogen-free conditions were maintained with food and water. This study of OVA induces inflammation and *in vivo* imaging was approved by the Animal Ethics Committee at the University of Gothenburg, Gothenburg, Sweden (permit no. 22-2016).

### Generation of MSCs with GFP and luciferase

Stable enhanced green fluorescent protein (e-GFP) and luciferase expressing cells were generated by infecting the MSCs with Lenti-virus that was generated in HEK-293T cell cotransfection of lentiviral vector pHAGE PGK-GFP-IRES-LUC-W (a gift from Darrell Kotton Addgene plasmid # 46793) with 4 expression vectors encoding the packaging proteins Gag-Pol, Rev, Tat, and the G protein of the vesicular stomatitis virus (VSV-G). With this protocol we obtained 5×10^8^ to 5×10^9^/ml of viral titers.

### Airway inflammation, in vivo imaging and broncho-alveolar lavage fluids (BAL) collection

An ovalbumin (OVA) mouse model of lung inflammation (Supplementary Fig. 10) was used to evaluate the migration potential of EV-treated MSCs *in vivo*. Intra-peritoneal (i.p) injection of OVA (8 μg/mice) were performed to sensitized mouse on day 1. On three consecutive days (14, 15 and 16 day) the mouse was intra-nasally (i.n) exposed to 100 μg/mice OVA (OVA / OVA group) or with PBS (OVA/PBS group). On day 17 post-sensitization mice from each group received 0.5 Million MSCs (expressing constitutive Luciferase and eGFP) that had either been incubated or not incubated with EVs for 48 hours. After 10 min, 30 min, 60 min, and 120 min, bioluminescence (photons/sec/cm^2^) from whole body of the mice was acquired with an IVIS spectrum (Caliper Life Sciences, Waltham, MA, USA) and analyzed by Living Image software (Version 4.0). Five minutes prior to imaging, i.p injection of D-Luciferin was administered to the mouse. Broncho alveolar lavage (BAL) from lung was collected 72 hours after the final OVA exposure as described previously (*55*).

### Isolation of EVs

EVs were isolated from conditioned cell medium by differential centrifugation with a filtration step. Briefly, 3-4 days culture medium was centrifuged at 300 × g for 10 minutes to remove cells. The supernatant was further centrifuged at 16,500 × g for 20 minutes. Then the supernatant was centrifugation at 120,000 × g for 3 hours (Type 45 Ti rotor, Beckman Coulter). The pellet was dissolved in PBS and protein concentration was measured by using BCA Protein assay kit (Pierce™, Thermo Fisher Scientific, Waltham, MA, USA).

### Reverse migration and invasion assay

The migration capacity and invasiveness of MSCs were evaluated using a 48-well Boyden chamber (Neuroprobe, Gaithersburg, MD, USA). In some experiments, MSCs were pre-incubated with mast cell-derived EVs for 48 hours before seeding and referred to as EV-treated MSCs. Five thousand cells/well were seeded to the bottom compartment and was separated from the upper chamber by a polycarbonate membrane with 8 µm pores. The membrane was pre-coated with 0.1% gelatin or 200 μg/ml ECM Gel from Engelbreth-Holm-Swarm murine sarcoma (Sigma-Aldrich). After being seeded, cells were allowed to adhere onto the membrane by inverting the chamber assembly upside down for 3.5 hours. Later the chamber was placed in the correct orientation and FBS was added in the upper compartment. After incubation for 12 hour at 37 °C, the membrane was removed and cells on the migrated sides were fixed in methanol (10 min), and stained with Giemsa (Histolab, Västra Frölunda, Sweden) for 1 h. Cell from the non-migrated side were wiped out before imaging. Three fields at 40 × magnifications were imaged. For the migratory inhibition experiments, MSCs were incubated with 100 nM of LY2157299 (Selleckchem, Munich, Germany) against TGFβ type-1 receptor. Each analysis was performed in triplicate.

### Scratch assay

Human MSCs were grown to 70-80 % confluence in 6-well plates and the monolayer cells were scratched with a 1 ml pipette tip across the center of the wells. After the cells had been washed with PBS, MEM plain medium with or without HMC-1-derived EVs (100 μg/ml) was added to the MSCs. Migratory cells from the scratched boundary were imaged after 24 and 48 hours.

### Gelatin zymography

The supernatant from MSCs, cultured with or without mast cell-derived EVs, were collected at 24 and 48 hour and electrophoresed onto zymogram precasted gels containing 10% gelatin (BioRad Laboratories, Hercules, CA, USA) with 5 × non-reducing loading buffers (Sigma Aldrich). Gels were renatured with 2.5% Triton X-100 (Sigma Aldrich) for 1 hour at room temperature and then incubated in development solution (50 mM Tris (pH 7.4), 5 mM CaCl_2_, 200 mM NaCl) at 37°C overnight. Gels were stained (coomassie brilliant blue) and destained (30% methanol and 10% acetic acid) until the clear bands appeared. Finally, the gel was incubated with stop solution (2 % acetic acid). Bands intensity was quantified by using ImageJ software.

### Sample digestion and nanoLC-MS analysis of identify membrane proteins on EVs

Proteomic analyses were performed at The Proteomics Core Facility at the Sahlgrenska Academy, University of Gothenburg. The samples in approximately 100 μl PBS were lysed by addition of Sodium dodecyl sulfate (SDS) to a final concentration of SDS 2% and 50 mM Triethylammoinium bicarbonate (TEAB). Total protein concentration was determined with Pierce™ BCA Protein Assay (Thermo Scientific). Aliquots containing 50 μg of each sample were digested with trypsin using the filter-aided sample preparation (FASP) method (*56*). Briefly, protein samples were reduced with 100 mM dithiothreitol at 60°C for 30 min, transferred to 30 kDa MWCO Pall Nanosep centrifugal filters (Sigma-Aldrich), washed with 8M urea solution repeatedly and alkylated by addition of methyl methanethiosulfonate to a final concentration of 10 mM. Digestion was performed in 50 mM TEAB, 1% sodium deoxycholate (SDC) over night at 37°C after addition of Trypsin (Pierce Trypsin Protease, MS Grade, Thermo Fisher Scientific) in a ratio of 1: 100 relative to protein amount. An additional portion of trypsin were added and incubated for another two hours followed by collection of the peptides by centrifugation. Samples were acidified to pH 2 by addition of TFA to precipitate SDC.

Samples were desalted using PepClean C18 spin columns (Thermo Fisher Scientific) according to the manufacturer’s guidelines prior to analysis on a Q Exactive mass spectrometer (Thermo Fisher Scientific) interfaced with Easy nLC 1000 liquid chromatography system. Peptides were separated using an in-house constructed C18 analytical column (200 × 0.075 mm I.D., 3 μm, Dr. Maisch, Germany) using the gradient from 5% to 25% acetonitrile in 0.1% formic acid for 75 min and finally from 25% to 80% acetonitrile in 0.1% formic acid for 5 min at a flow of 200 nL/min. Precursor ion mass spectra were recorded in positive ion mode at a resolution of 70 000 and a mass range of *m/z* 400 to 1600. The 10 most intense precursor ions were fragmented using HCD at a collision energy of 27, and MS/MS spectra were recorded in a scan range of *m/z* 200 to 2000 and a resolution of 35 000. Charge states 2 to 6 were selected for fragmentation, and dynamic exclusion was set to 30 s. Samples were re-analyzed with exclusion lists of *m/z* values of the identified peptides at 1% FDR with a 10 min retention time generated after database searching (as described below) of previous LCMS runs.

Data analysis was performed utilizing Proteome Discoverer version 1.4 (Thermo Fisher Scientific) against Human Swissprot Database version March 2015 (Swiss Institute of Bioinformatics, Switzerland). Mascot 2.3.2.0 (Matrix Science) was used as a search engine with precursor mass tolerance of 5 ppm and fragment mass tolerance of 100 mmu. Tryptic peptides were accepted with one missed cleavage, methionine oxidation was set as variable modifications and cysteine alkylation as static modification. The detected peptide threshold in the software was 1% False Discovery Rate by searching against a reversed database and identified proteins were grouped by sharing the same sequences to minimize redundancy.

### Immunofluorescence microscopy

Isolated lungs were perfused with O.C.T compound (Tissue Tek, Alphen aan den Rijn, The Netherlands) and rapidly freezed in liquid nitrogen (LN_2_) and 40µm sections were made on positively charged slides. To localize immunofluorescence signal from mouse lung, sections were mounted and fixed with 3.7% paraformaldehyde at RT for 10 min, permeabilized for 5 min with 0.2% triton X-100, washed and blocked for 1 hour in 3% BSA. Sudan black (0.1% in 70% EtOH) staining was performed for 20 minutes followed by PBS three washes to remove the auto-fluorescence. Finally, incubation with GFP-AF-488 antibody (1:200, A21311, Life Technology) was performed for 1 hour at RT. After three PBS washes, the cells were further stained with DAPI (Sigma Aldrich) and cover slips were mounted using Gold anti-fade mounting reagent (Invitrogen, Carlsbad, CA,USA) and observed under fluorescence light microscope (Axio Observer, Zeiss, Oberkochen, Germany). Above-mentioned staining protocol was also performed to stain MSCs for nuclear SMAD2 (s-20, Sc-6200, Santa Cruz Biotechnology, CA, USA) after EV-treatment, with the exception that the Sudan black staining was not included. Nuclear expression of SMAD2 was evaluated with Velocity Image analysis software (PerkinElmer, Chicago, IL, USA).

### Transmission electron microscopy

To describe the nano-structures in EVs derived from primary human mast cells, we performed transmission electron microscopy (TEM). Isolated EVs described in section “Isolation of EVs”, were bottom loaded and floated on optiprep density gradient (0, 20, 22, 24, 26, 28, 30 and 50%) and centrifuged at 182,300 × g for 16 hours (SW40-Ti Rotor) to separate them from free proteins. Nine different optiprep fractions were collected and samples from fraction no.2 was subjected for TEM as described in our previous study.

### Labeling of EVs and uptake

EVs was obtained as described in “Isolation of EVs” section and were labeled with PKH67 Green Fluorescent Cell Linker Kit (Sigma Aldrich) as per manufacturer’s protocol and as previously (*57*) with modifications in the removal of unbound dyes. The labeled EVs were loaded at the bottom of an OptiPrep cushion (0, 20, 30 and 50%) and centrifuged at 182,300 × g for 4 hours (SW40-Ti Rotor) to separate them from free unbound dye. The lipid labeled EVs were collected from the interphase between 20-30 % and washed in PBS for 120,000 × g for 3.5 hours (Type 45 Ti rotor, Beckman Coulter). Washed EVs were incubated with the MSCs (4000 cell/cm^2^) for 4 or 16 hours at 4 °C and 37°C. FACS was performed on the cells to determine the uptake rate. For visualization, EVs were incubated with MSCs for 4 hour, paraformaldehyde (3%) fixed, DAPI stained and imaged under a fluorescence light microscope (Axio Observer, Zeiss).

### Fluorescence Correlation Spectroscopy

Freshly isolated EVs from HMC1.2 cells were labeled either alone with optimized concentration of DiO (Life Technologies, Thermo Fisher Scientific) lipid dye (0.22 µg/ml), TGFβ1-Alexa647 (1:50) and CD63-PE (1:20) or in combination of two labels (DiO/TGFβ1-AF647 or TGFβ1-AF647/CD63-PE or DiO/CD63-PE). All labeled EVs were purified from free label using optiprep density cushion described in the “Labeling of EVs and uptake” section. Washed pellets were analyzed by dual color Fluorescence Cross-Correlation Spectroscopy (FCCS). Two different FCS/FCCS setups were used: A confocal microscope (FCS-equipped Zeiss 780) fully equipped for FCS and FCCS measurements. On this setup, we used mainly the 488 nm and 633 nm laser lines, but to some extent also the 514 nm and the 561 nm laser lines. The 488 nm laser line resulted in a focus radius ω_0_=0.25 μm and a volume of 0.45 femtoliters (fL), while the 633 nm laser line gave a focus with ω_0_=0.29 μm and a volume of 0.65 fL. Analysis of the FCS/FCCS curves was performed using the Zeiss Zen software. In addition, a home-built FCCS setup based on a 488 nm line (Argon laser, Lasos GmbH) and a 594 nm line (HeNe laser, Laser2000 GmbH) was used. In this setup the 488 focus had ω_0_=0.36 μm and a volume of 1.5 fL, while the 594 focus had ω_0_=0.39 μm and a volume of 2.4 fL. Emission filters ET535/70 and ET700/75 (Chroma) were used. The correlator was ALV-5000 (ALV GmbH). Analysis was performed using the ALV-5000 software and Origin 9.1 (Originlab Corporation, USA).

### TGFβ-1 detection on EVs

Amount of TGFβ-1 in the supernatant of MSCs and on the HMC1 cell line derived EVs were performed using a TGFβ-1 ELISA Ready-SET-Go kit (eBioscience, Affymetrix) according to the instruction of the manufacturer. To measure the relative level of TGFβ-1 on primary human matured mast cells on captured CD63^+^ and TGFβ-1^+^ EVs we used high sensitive chemiluminescence based detection system (Roche Molecular Systems, In, USA). For this experiment we used CD63 antibody to coat the ELISA plate and remaining antibodies were used was from TGFβ-1 ELISA Ready-SET-Go kit.

### Quantitative real time polymerase chain reaction

Total RNA was isolated from MSCs using miRCURY™ RNA isolation kit for cell and clant (Exiqon, Vedbaek, Denmark). TURBO™ DNase treatment and removal reagents (Ambion, Life Technologies) was used to remove contaminating DNA from RNA preparation and the concentration and purity of RNA were evaluated by NanoDrop (Thermo Scientific). cDNA was synthesized from 200 ng total RNA by using iScript™ cDNA Synthesis Kit (BioRad) according to manufacturer’s protocol. Quantitative real time PCR was performed with SsoAdvanced™ Universal SYBR^®^ Green Supermix on BioRad CFX96™ system. cDNA was denatured for 30 seconds at 95°C, and then 95°C for 15 seconds, annealing at 60°C for 30 seconds for 40 cycles. The primers were obtained from Sigma (KiCqStart^®^ primers): TGFB1, SMAD2, MMP2 and EF1. Data was collected by software and analyzed. EF1 was used to normalize the data. 2_T_^−ΔΔC^ method was used to determine relative changes in gene expression.

### Western blotting

MSCs were lysed in RIPA buffer (Cell Signaling Technology ST, Danvers, MA, USA), 1 mM PMSF, 1 mM Na_3_VO_4_, and 1 mM NaF. Twenty micrograms of protein lysate were subjected to SDS-PAGE and transferred onto Nitrocellulose membranes. Membranes were blocked with 5% BSA in TBS containing 0.05% Tween-20 and incubated with primary antibodies at 4°C overnight and HRP conjugated secondary antibodies (1:10000, NA931V, NA9340, NA9310V, GE Healthcare) for 1 hour at room temperature. The proteins were detected with ECL Prime Western Detection (GE Healthcare) according to the manufacturer’s protocol. The antibodies used were as following: β-actin (1:10,000, 13E5, #4970S, Cell Signaling Technology), TSG101 (1:1000, ab83, Abcam), CD81 (1:1000, sc9158, Santa Cruz Biotechnology), EEA1 (1:1000, sc33585, Santa Cruz Biotechnology), LAMP1(1:1000, ab24170, Abcam), FLAG-antibody (1:5000 #M2, F3165, Sigma), and pSMAD2 (1:500; clone; Santa Cruz Biotechnology).

### Particle number and size measurements

The size and concentration of EVs were measured using ZetaView^®^ PMX110 (Particle Metrix, GmbH, Meerbusch, Germany). Each EV sample were diluted in PBS in a range of 1:1000~1:5000 and were injected into the instrument. The chamber temperature was maintained automatically. Measurements were obtained in triplicate and each individual data was obtained from two stationary layers with five times measurement in each layer. Sensitivity of camera was fixed at 70 in all measurements. Data were analyzed using ZetaView analysis software version 8.2.30.1 with a minimum size of 5 nm, a maximum size of 1000 nm, and a minimum brightness of 20.

### Lineage differentiation of MSCs

MSCs treated with 100 μg/ml EVs for 48 hours were cultured in lineage differentiation medium for 15 days as per manufacturer instruction (Human mesenchymal stem cell functional identification kit, SC006, R&D Systems). Lineage specific marker for Adipocytes (FABP4, Oil red O) and Osteocytes (Osteocalcin, Alizarin Red) was probed and analyzed in respective cells as described in previous published guidelines (*58*).

### TGFβ-1 knockdown

Stable doxycycline-regulated Cas-9-expressing human mast cell (HMC-1.2) was generated by cloning pCW-Cas9 containing plasmid (Gift from Eric Lander and David Sabatini; Addgene Plasmid #52961). Target gRNA (ACCAAAGCAGGGTTCACTAC) against human TGFbeta-1 gene was cloned in plasmid backbone of pLX-sgRNA (Gift from Eric Lander and David Sabatini (Addgene Plasmid #50662) with Reverse (R1) primer 5’-GTAGTGAACCCTGCTTTGGTCGGTGTTTCGTCCTTTCC-3’ and Forwards primers (F2) 5’-GACCAAAGCAGGGTTCACTACGTTTTAGAGCTAGAAATAGCAA-3’. The gRNA was chosen from the library B of Human GeCKO v2 Library (2- Plasmid System - lentiGuide-Puro) (cat #1000000049). The pLX-TGFB1 gRNA carrying plasmid was expanded and purified from chemical competent *E.coli* (MAX Efficiency® Stbl2, Life Technology) and transfected to Cas-9-HMC1.2 with doxycycline (0.5 μg/ml). Knockdown cloned were selected using limiting dilution of cells under blasticidin. Efficiency of knockdown was tested by estimating the level of TGFβ-1 protein in knockdown cells lysate (Supplementary Fig. 5).

### Localization of EVs and isolation of organelles

Isolated EVs (as described in method section of “Isolation of EVs”) were loaded at the bottom of optiprep gradient (0, 20, 22, 24, 26, 28, 35 and 50 %) and centrifuged at 182,300 × g for 16 hours (SW40-Ti Rotor) to purify the EVs from layer 20-22 % of the optiprep gradient. The surface of the optiprep purified EVs (10 μg/ml) were biotinylated by incubating it with EZ-Link Sulfo-NHS-Biotin (Thermo Scientific) as per the manufacturer recommended. Free-biotin was removed from biotinylated-EVs via dialysis by sequentially changing PBS after 2 hours (room temperature), overnight (4°C) and finally for 2 hours (RT). These biotinylated-EVs were later incubated with HEK-293T cells for 60 mins and organelles were isolated from these cells using lysosome enrichment kit (Thermo Scientific) with manufacturer’s guideline with further modification to enrich lysosomes and endosomal compartments from the organelle pool. Briefly, 100 mg HEK293-T were mixed with reagent “A” with short vortex followed by lysis via sonication pulse (25-30 burst, 2 min each, 4°C). This suspension was mixed uniformly with equal volume of reagents “B” and centrifuged (500 ×g, 10 mins 4°C) to collect the supernatant. This supernatant was loaded onto the top of an iodixanol density gradient (17, 20, 23, 27, and 30%) and centrifuged at 145,000 × g for 2 hours at 4°C. After first round of ultracentrifugation, first two fractions (green and red, Supplementary Fig. 9a) were collected and bottom loaded on two separate tubes with different iodixanol density (18, 20, 22, 22, 24, 26, 28, 30, 35, and 50 %) for second round of ultracentrifugation (182,300 × g, SW40-Ti Rotor, 16 hours, 4°C). Later the fractions were collected from top to bottom (1 ml each for 10 factions) and subjected for the western blotting analysis of lysosomes (LAMP1) and endosomes (EEA1) enrichment markers to defining the distribution of density. From these fractions, lysosomes enriched fraction (LEF; fraction 2-3) and endosomes enriched fraction (EEF; fraction 7-9) were used to locate the biotinylated-EVs (Supplementary Fig. 9b).

In order to locate the biotinylated-EVs in these fractions (LEF and EEF) two complementary approach were used. Firstly, LEF and EEF were both incubated separately with streptavidin-coated beads and probed for total bounded protein or EEA1 and LAMP1 antibody (Fig. 4C). In second approach, LEF and EEF fractions were both incubated separately either with EEA1 or LAMP1-coated beads and probed to detect biotinylated proteins in respective fraction.

## Legends for Supplementary Data

**Supplementary Table 1:** List of proteins obtained from membrane proteomics from mast cells derived EVs with the number of peptide hits.

**Supplementary Figure 1:**
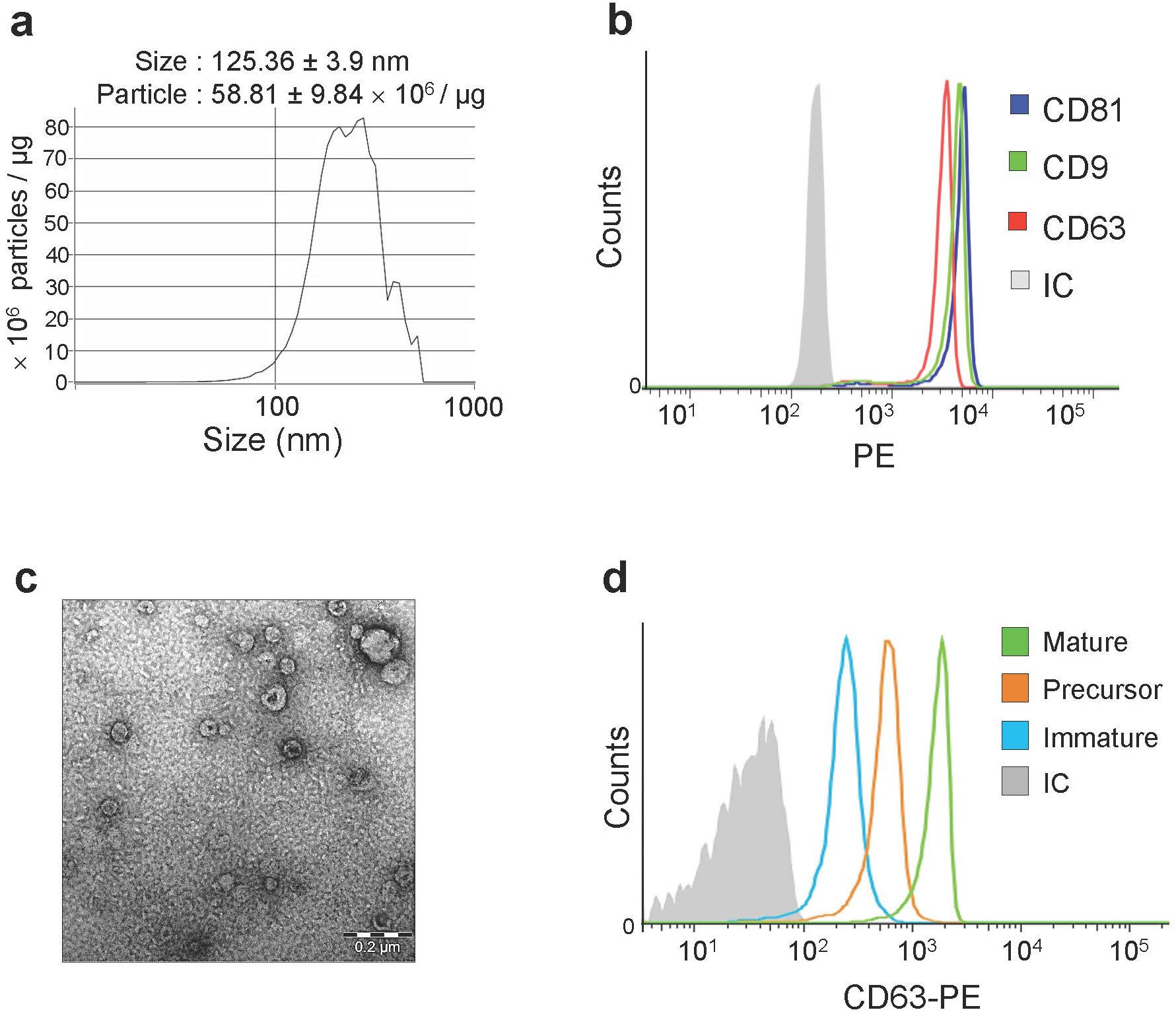
Characterization of mast cell-derived extracellular vesicles. **(a)** Size distribution and particle concentration of mast cell-derived EVs per unit of protein using nanoparticle-tracking analysis. **(b)** EVs were captured on anti-CD63-coated beads, and the presence of CD63, CD81, and CD9 was determined with flow cytometry. **(c)** Electron microscopy was performed on the EVs from differentiated matured human mast cells (week 6-8). **(d)** EVs isolated from primary human mast cell were incubated with CD63 antibody coated magnetic beads and FACS analysis was made to determine the levels of CD63 on those EVs.

**Supplementary Figure 2:**
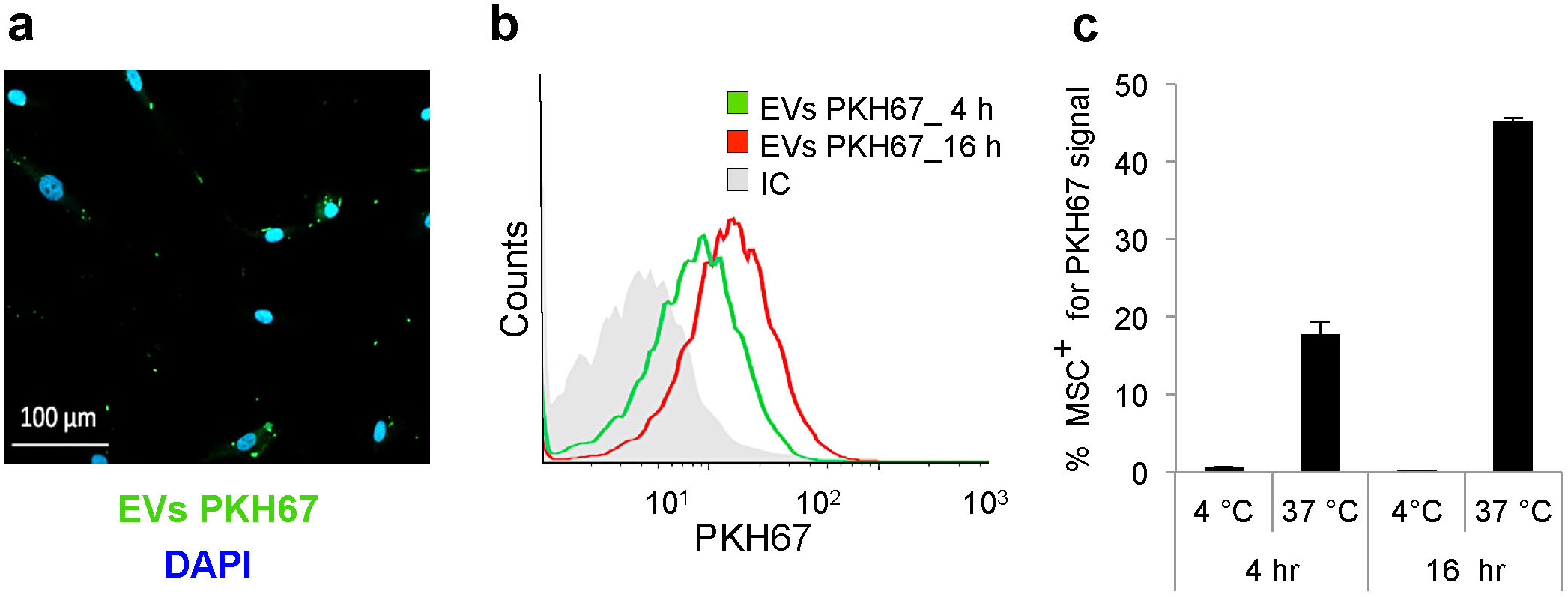
EV uptake by MSCs is an active process. **(a)** Immunofluorescence image of MSCs that had been taking up PKH67-labeled EVs after 4 hours of co-incubation (scale bar is 100 μm). (**b**) Flow cytometry analysis of MSCs that were incubated with PKH67-labled EVs for 4 h or 16 h. **(c)** Percentage of MSCs that were positive for PKH67-labled EVs after incubation at 4°C or 37 °C for 4 h or 16 h.

**Supplementary Figure 3:**
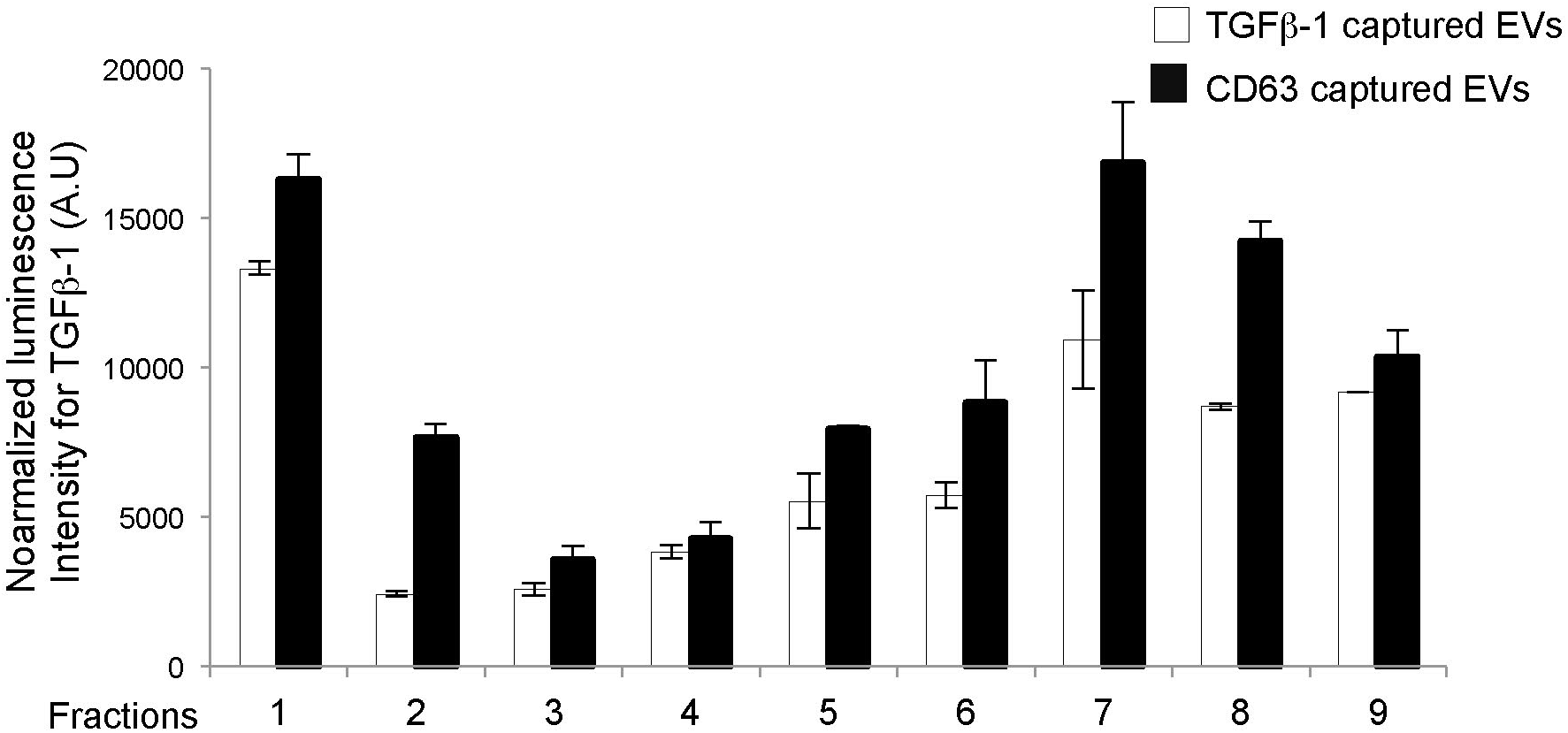
TGFβ-1 distribution on EVs derived from primary human mast cells. EVs from matured mast cells (week 6-8) were floated on optiprep density gradient and relative levels of TGFβ-1 were determined using sandwich ELISA for TGFβ-1 and CD63 captured EVs (n=3).

**Supplementary Figure 4:**
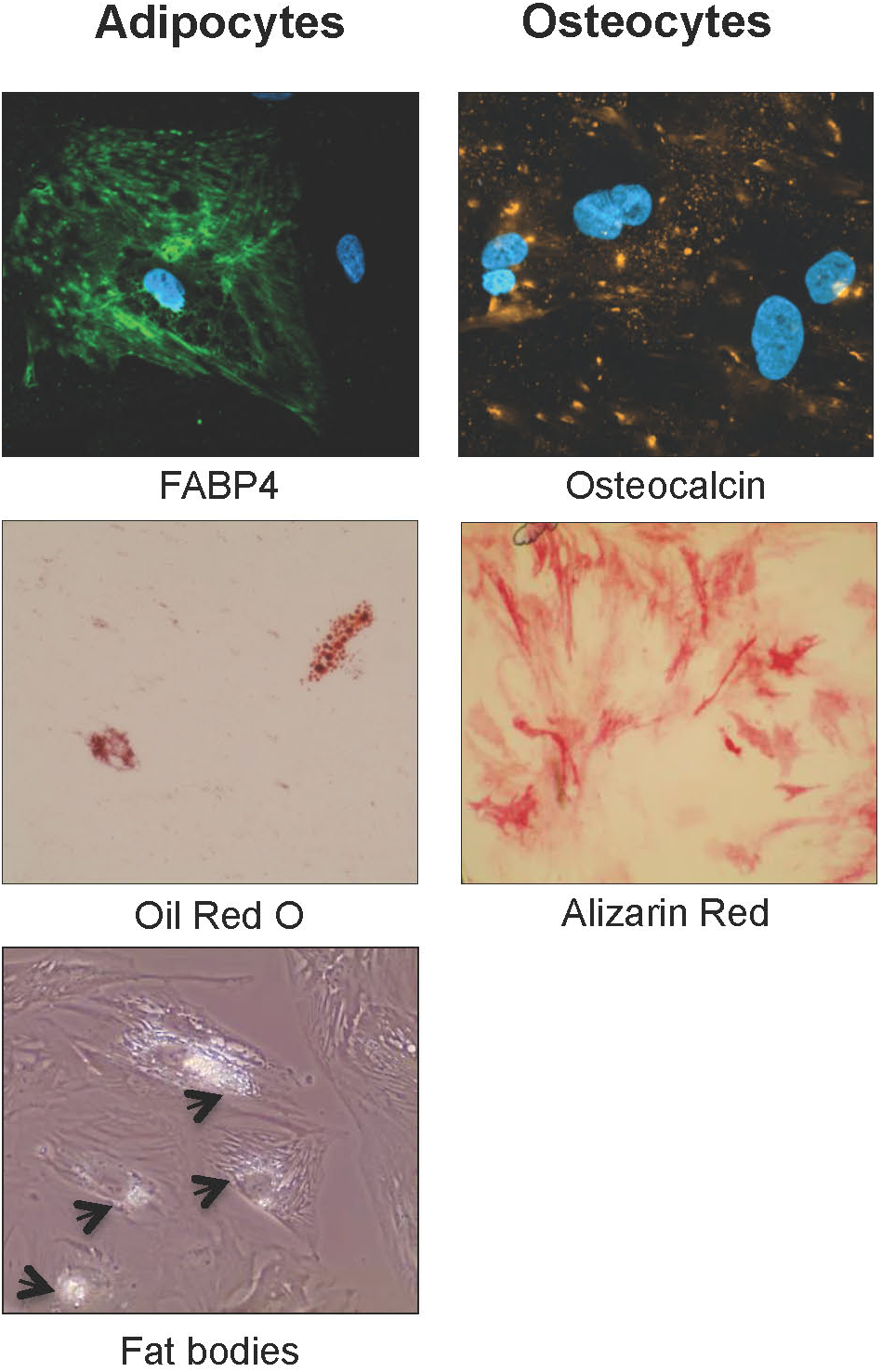
EVs do not alter the multipotency of MSCs. MSCs were incubated with mast cell-derived EVs for 48 h and then differentiated into the adipocytic, and osteocytic. After 15 days of culturing MSCs in corresponding differentiation medium, the cells were labeled with lineage-specific markers for **(a)** adipocytes (FABP4, Oil Red O and fat bodies), and **(b)** osteocytes (Osteocalcin and Alizarin Red).

**Supplementary Figure 5:**
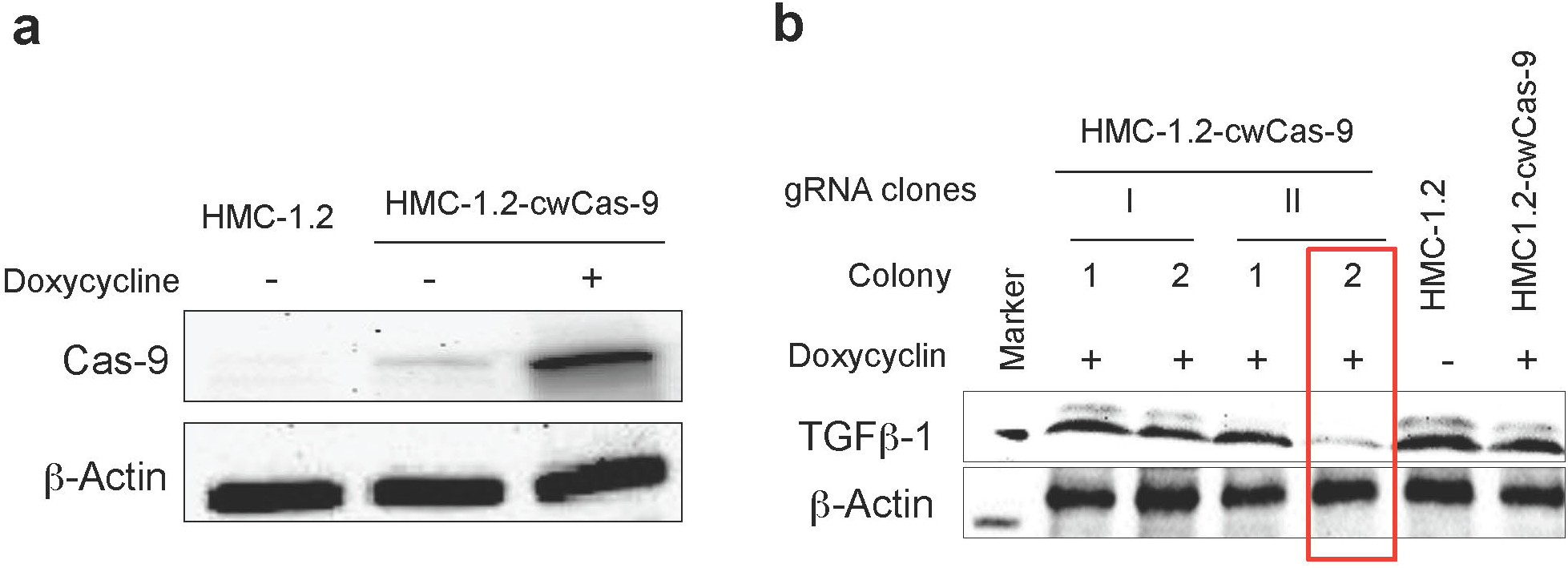
Doxycycline-regulatable Cas-9-expression system in mast cells. **(a)** Immunoblotting analysis of FLAG-tagged Cas-9 nuclease, regulated by doxycycline, in HMC-1.2 cells transduced with pCW-Cas9. **(b)** The efficiency of TGFβ-1 gRNA clones in Cas-9-expressing HMC1.2 cells was estimated by immunoblotting of TGFB-1 in various clones.

**Supplementary Figure 6:**
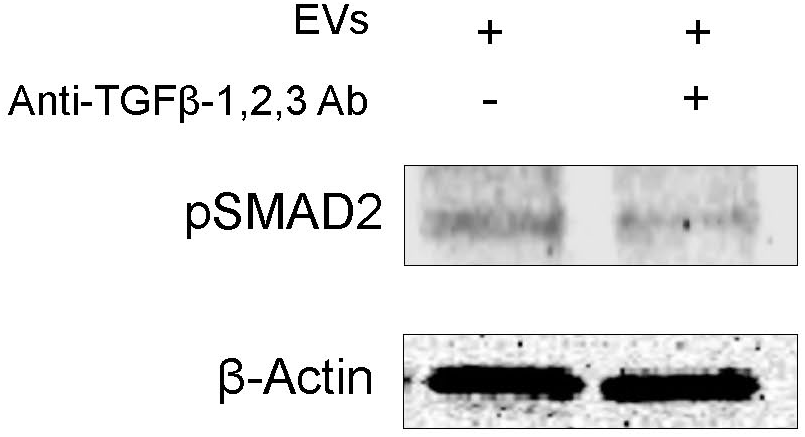
EV-associated TGFβ-1 regulates activation of TGFβ-1 signaling. Immunoblot analysis of pSMAD in MSCs 60 minutes after treatment with EV-bound TGFβ-1 and after blocking the EV-bound TGFβ-1 using blocking antibody.

**Supplementary Figure 7:**
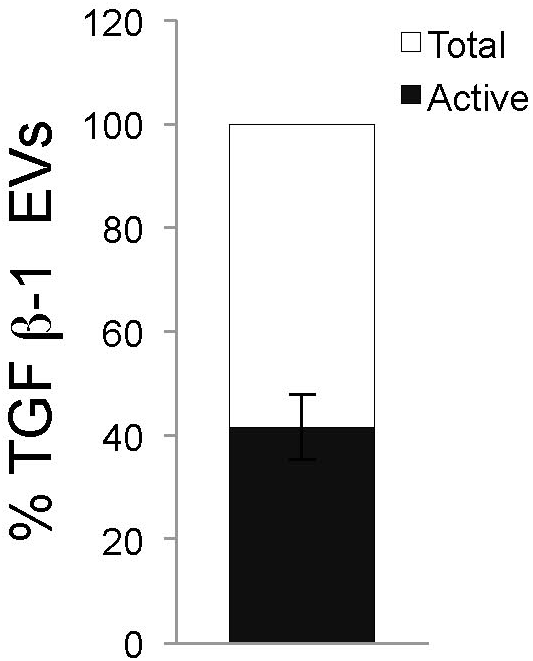
Contribution of TGFβ-1 in total EVs. Contribution of TGFβ-1 in active and total form was evaluated using ELISA and presented as percentage (n = 3).

**Supplementary Figure 8:**
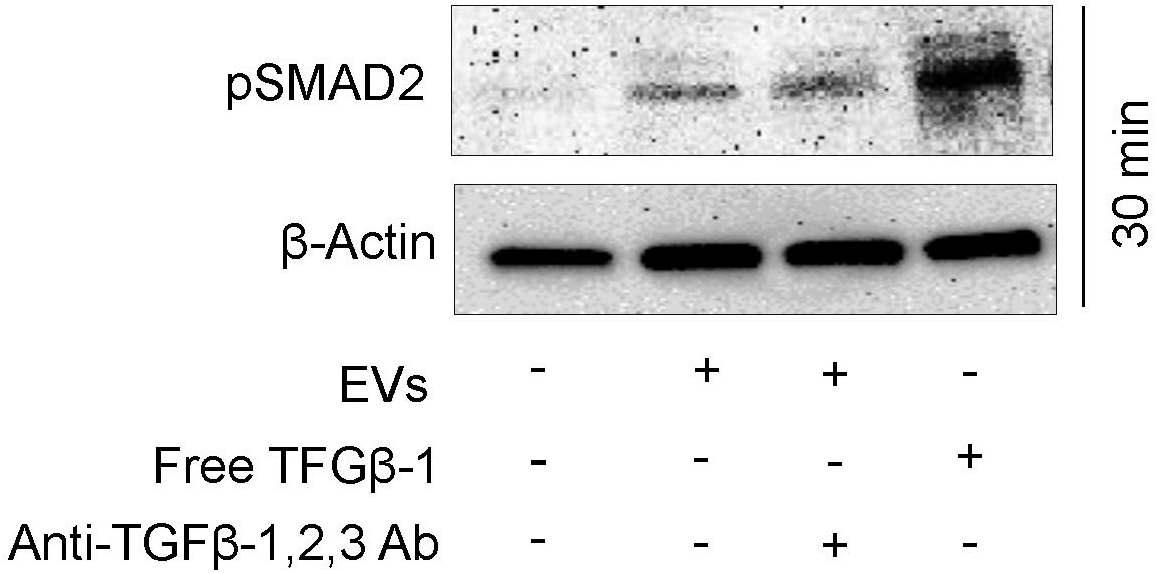
EV-associated TGFβ-1 regulates activation of TGFβ-1 signaling. Immunoblot analysis of pSMAD in MSCs after 30 minutes treatment with free TGFβ-1 or EV, as well as after blocking the EV-bound TGFβ-1.

**Supplementary Figure 9:**
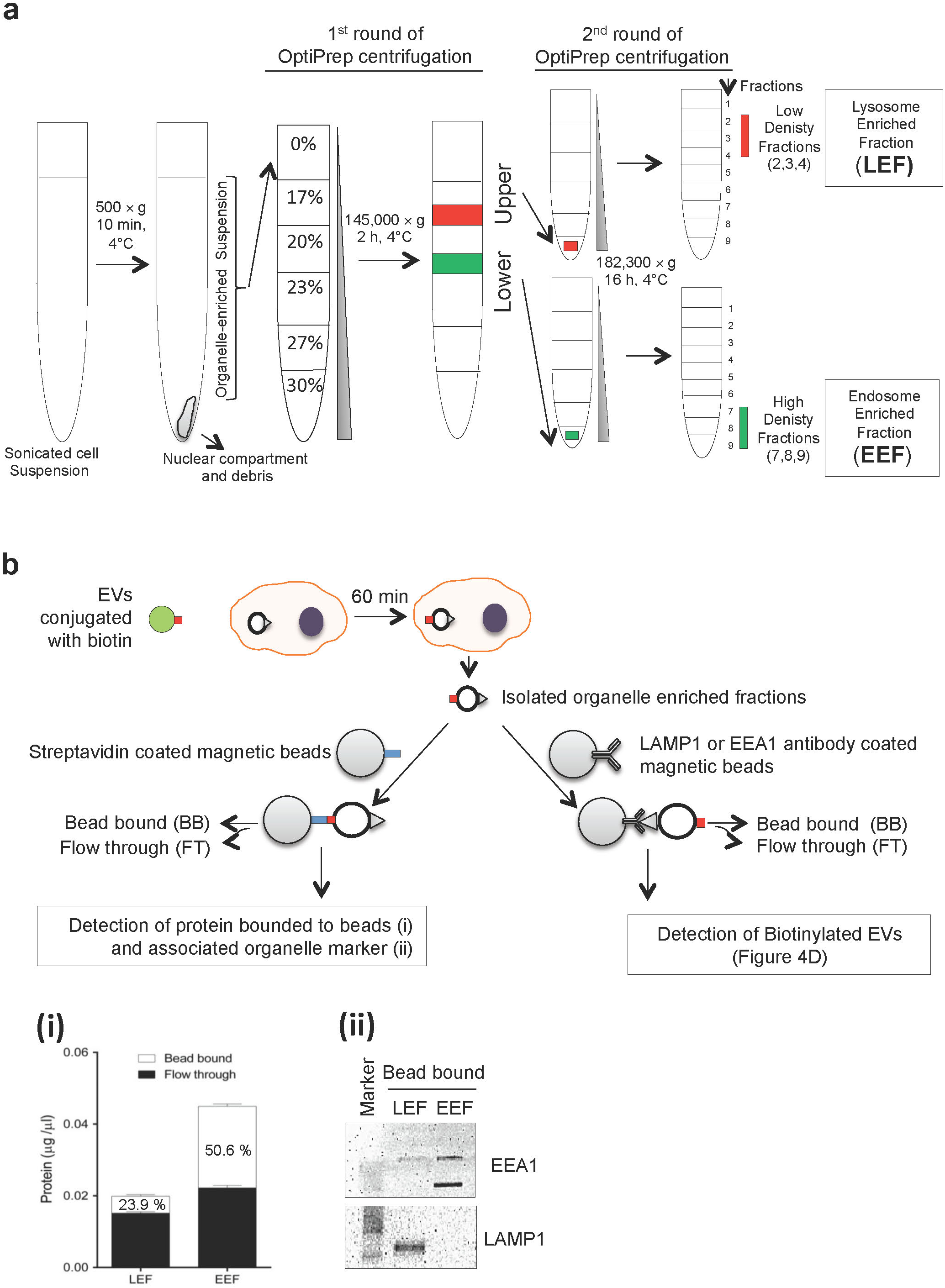
Outline of the method to isolate the lysosome-enriched and endosome-enriched compartments. **(a)** Outline for separating/enriching endosomes and lysosomes. A total of 100 mg of cells was treated with biotinylated EVs (100 μg/ml) for 60 min, after which the cells were lysed and centrifuged to remove cells and debris. Supernatants were top loaded and floated on an iodixanol density gradient followed by a second round of iodixanol density gradient separation of the upper and lower fractions obtained from the first round of iodixanol density gradient. The final fractions for the endosome-enriched fraction (EEF) and the lysosome-enriched fraction (LEF) were collected for subsequent experiments. **(b)** Biotinylated EVs were incubated with HEK293-T cells for 60 min, and organelles were isolated as described in Supplementary Fig. 9a. Isolated organelles were incubated independently with streptavidin-coated magnetic beads or organelle-specific (LAMP1 or EEA1) antibody-coated magnetic beads (left side). **(i)** Total protein in the bead bound (BB) and flow through fractions was measured to evaluate the biotinylated protein in the lysosome-enriched fraction (LEF) and endosome-enriched fraction (EEF). **(ii)** Organelle-specific markers (LAMP1 or EEA1) were evaluated by immunoblotting to detect their presence in the LEF and the EEF that had bound to the streptavidin-coated beads. In parallel (right side), the presence of biotinylated protein in organelles isolated from magnetic beads coated with LAMP1 or EEA1 antibodies was evaluated by immunoblotting (Fig. 4d) with a streptavidin antibody.

**Supplementary Figure 10:**
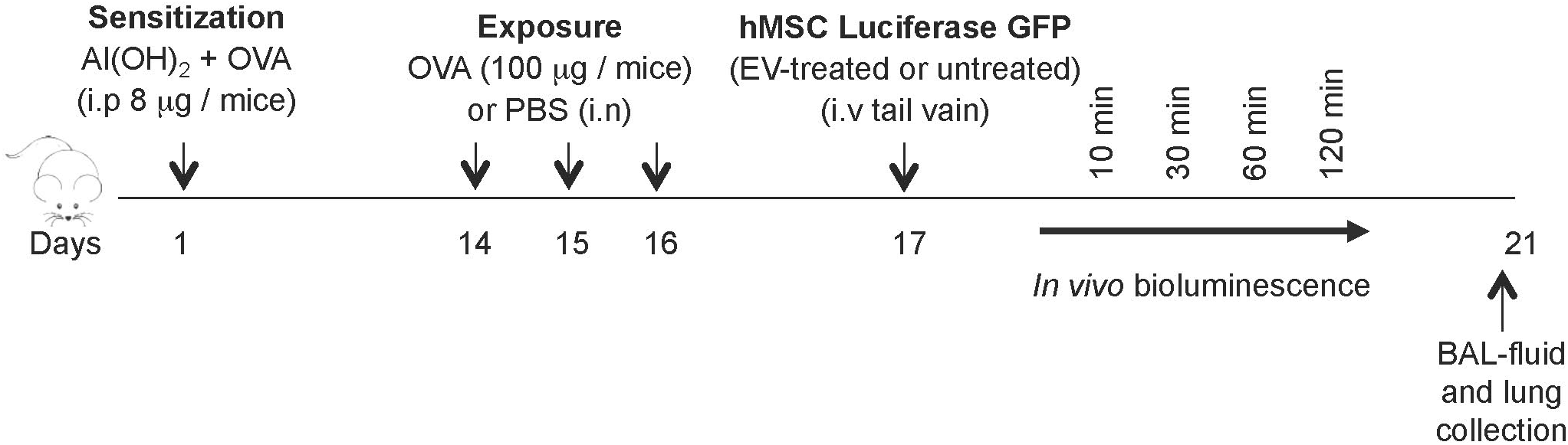
Standard protocol for OVA-induced airway inflammation in a mouse model.

**Supplementary Figure 11:**
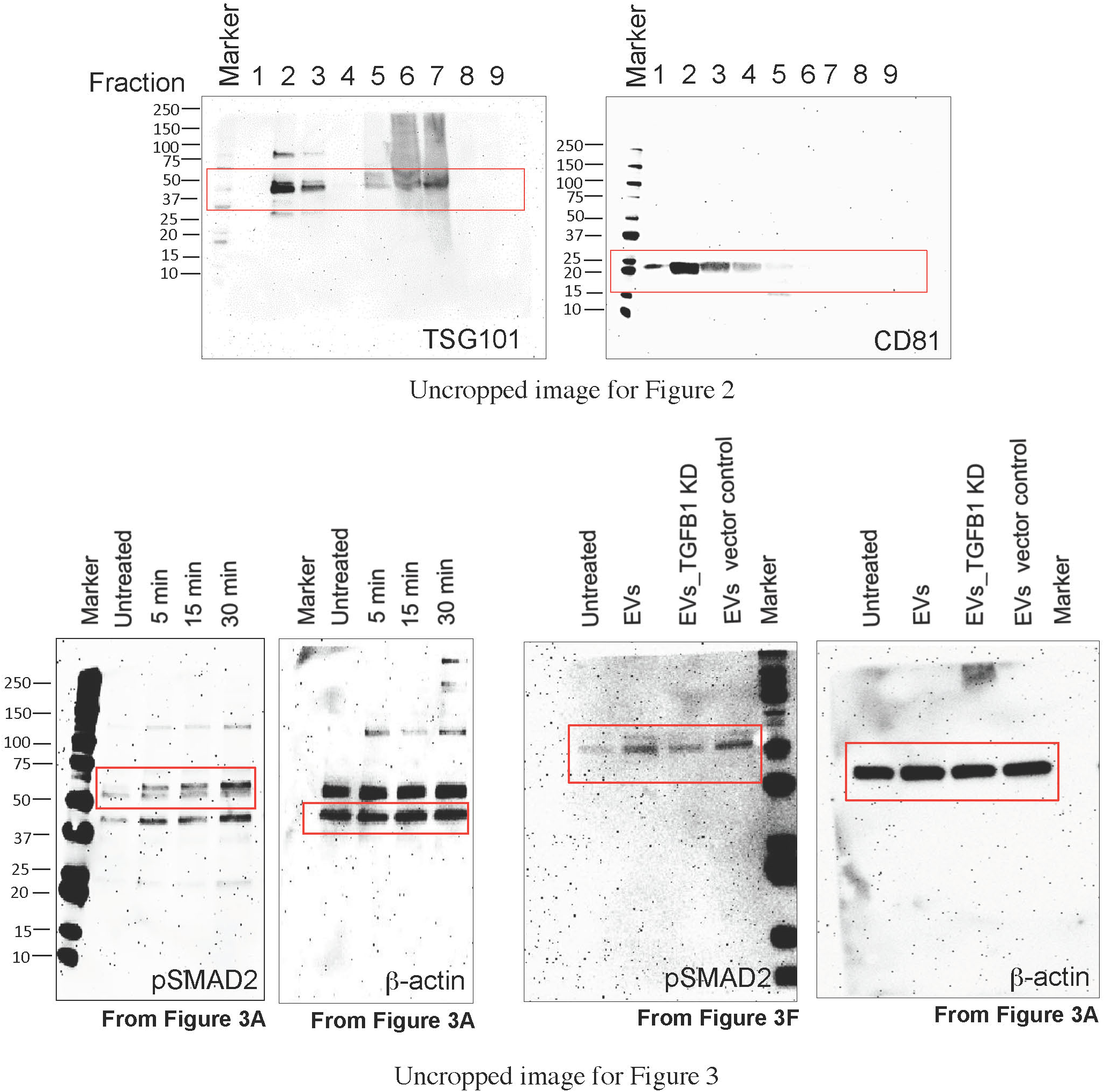
Uncropped western blot images for figure 2 and figure 3.

**Supplementary Figure 12:**
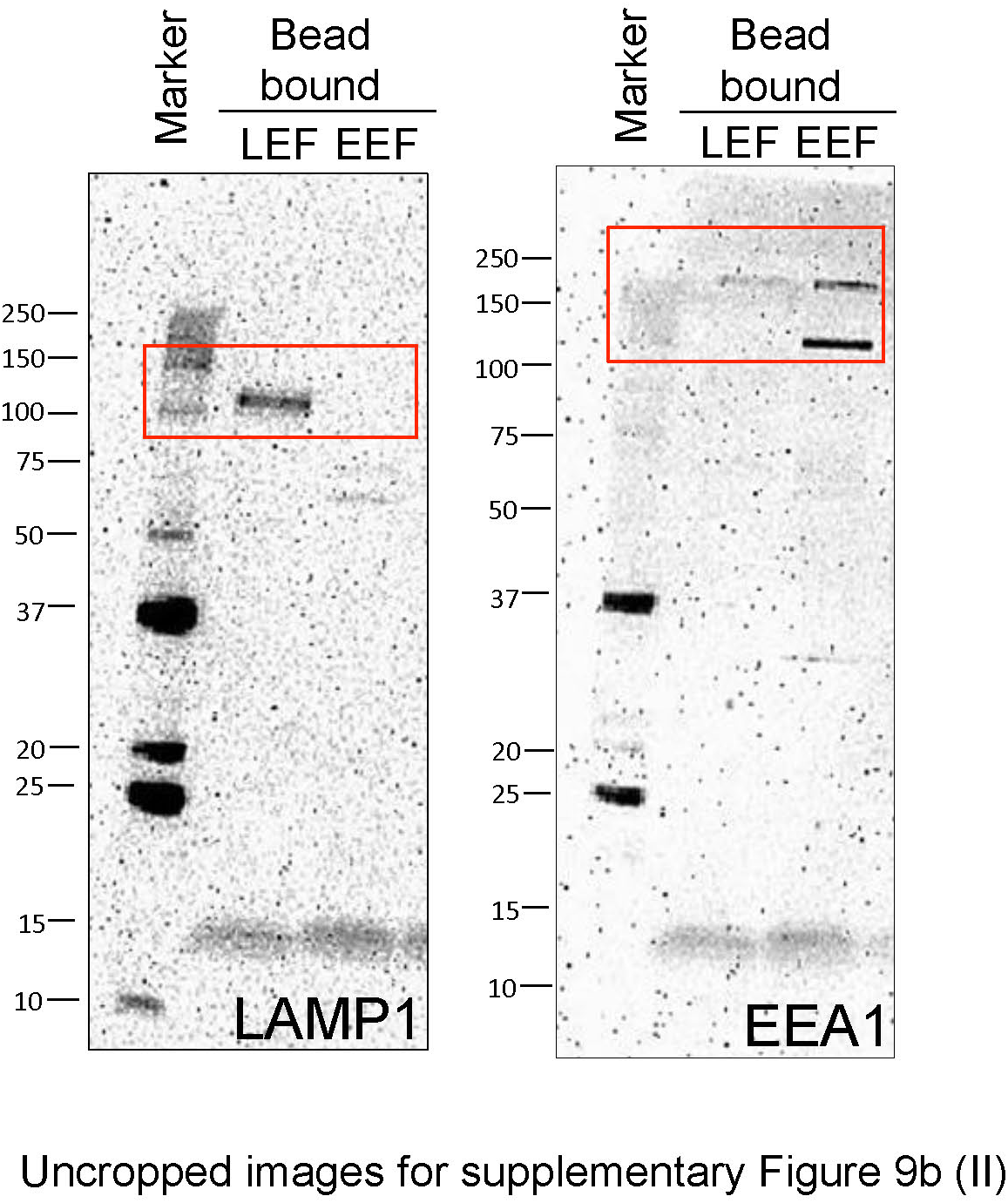
Uncropped western blot images for supplementary figure 9b.

## ACKNOWLEDGEMENT

We thank the Proteomics Core Facility at Sahlgrenska Academy, Gothenburg University for performing the proteomic analysis. We thank Madeleine Rådinger (Krefting Research Centre, University of Gothenburg, Sweden) for kind support for providing valuable input in usage of animal model and ethical application. We thank Joydeep Bhadury for his technical guidance and sharing resources to develop CRISPR/Cas-9 cells. Dr. Lisa Nilsson (Sahlgrenska Cancer Center University of Gothenburg, Sweden) for mouse handling and tail vein injections. The intramural finding from Herman Krefting Foundation for Allergy and Asthma Research, Swedish Cancer Foundation, Swedish Research Council and Swedish Heart and Lung Foundation VBG-group Centre for Allergy and Asthma research to supported this work. GS is supported by European Academy of Allergy and Clinical Immunology, Assar Gabrielssons Foundation, Lundgren Foundation, Sahlgrenska University hospital and Sahlgrenska Academy.

